# An unrecognized and crucial role of chloroplast division in leaf variegation in *Arabidopsis thaliana*

**DOI:** 10.1101/2025.04.06.647415

**Authors:** Wenjuan Wu, Wei Guo, Haojie Zhu, Di Li, Zhiyi Zhang, Danni Lin, Meiying Qu, Zhenjia Yu, Jirong Huang

## Abstract

Leaf variegation provides a valuable model for elucidating the molecular mechanisms that underlie chloroplast biogenesis, including both chloroplast development and division. While there has been notable advancement in comprehending the role of the *Arabidopsis Yellow Variegated2* (*VAR2*), which encodes the AtFtsH2 subunit of the thylakoid FtsH metalloprotease complex, the exact mechanism governing leaf variegation formation remains elusive. In this study, time-course microscopy analyses of chloroplast number per cell from the first leaf pair revealed that most *var2* cells lack chloroplasts. Interestingly, chloroplast biogenesis in the green sector of *var2* leaves is significantly delayed but prolonged compared to the wild type (WT), leading to increased heterogeneity in chloroplast number and size among cells. These results suggest that chloroplast biogenesis in the green sector can partially compensate for the defect in the white sector. Additionally, approximately 15% of cells in *var2* white sectors are devoid of plastids, and chloroplast number in guard cells is significantly lower in *var2* than in WT, indicating that *VAR2* mutations impair chloroplast division. Consistently, the *var2* phenotype is exacerbated by knocking out key plastid division genes, including *Paralog of Accumulation and Replication of Chloroplasts6* (*PARC6*) and *Plastid Division1* (*PDV1*), but is fully rescued by overexpressing *PDV1* or *PDV2*. Furthermore, *VAR2* mutations inhibit etioplast development, while accelerating chloroplast biogenesis by knocking out *Constitutively Photomorphogenic1* (*COP1*) rescues the *var2* phenotype. Together, our findings reveal a novel role for VAR2 in plastid division and its essential function in chloroplast biogenesis, providing new insights into the mechanism underlying variegated leaf formation.

## Introduction

Chloroplast biogenesis in higher plants is a fundamental process that reconstructs photosynthesis and provides sufficient energy and materials for autotrophic growth. In general, chloroplast biogenesis is regulated by the two basic events, namely development and division of chloroplasts (Sakamoto *et al*., 2008). Chloroplast development is a light-triggered dynamic event facilitating any kind of non-photosynthetic plastids, such as small, undifferentiated proplastids in meristematic cells and differentiated plastids (etioplasts, chromoplasts and leucoplasts), into chloroplasts. It is well documented that chloroplast development is finely regulated by a series of important processes, such as coordinated expression of nuclear and plastidic genes, protein import into chloroplasts and subsequent targeting to the sub-organelle compartments, biogenesis of thylakoid membranes and photosynthetic complexes, etc (Jarvis and López-Juez, 2013). On the other hand, the number of semiautonomous chloroplasts in a cell is determined by its division through binary fission, which keeps pace with cell differentiation, proliferation and expansion. These two events of chloroplast biogenesis are closely linked but also independent, leading to dramatic variation in chloroplast number in different cell types (Fang *et al*., 2022; Pyke, 1997). For example, some *arc* (accumulation and replication of chloroplast) mutants exhibit an increased number of small chloroplasts, while some have only one or two large chloroplasts (Chen *et al*., 2018). This indicates that chloroplast number and size, which reflect the outcomes of chloroplast division and development, respectively, can mutually compensated for each other.

The variegated leaf is an idea phenotype to elucidate molecular mechanisms underlying chloroplast biogenesis, since green sectors with normal chloroplasts and yellow or white sectors with non-photosynthetic plastids coexist in the same genetic background, and severity of leaf variegation is significantly affected by growth conditions and developmental stages (Liu *et al*., 2010). To date, many variegated-leaf mutants, such as *immutans* (*im*), *chloroplast mutator* (*chm*), *thylakoid formation 1* (*thf1*), *variegated1* (*var1*) and *var2*, have been identified in the model plant *Arabidopsis thaliana* (Liu *et al*., 2010; Yu *et al*., 2007). Among them, *VAR1* and *VAR2*, which encode FtsH5 and FtsH2 subunits of the thylakoid FtsH metalloprotease complex, respectively, are the most extensively studied. *VAR1/FtsH5* and *FtsH1* are functionally redundant and are collectively referred to as type A, while *VAR2/FtsH2* and *FtsH8* are interchangeable and classified as type B. The two major subunits, FtsH2/VAR2 and FtsH5/VAR1, assemble with the minor subunits FtsH1 and FtsH8 to form the hetero-hexameric FtsH complex, which plays an important role in the degradation of the D1 protein at the reaction center of photosystem II (PSII) as well as in the removal of misfolded polypeptides and protein aggregates (Kato *et al*., 2018; Kato *et al*., 2009). Several hypotheses have been proposed to explain form the formation of leaf variegation, one of which is a threshold model. This model suggests that a certain level of FtsH activity is required for chloroplast development and green sector formation (Aluru *et al*., 2006). Genetic screening for second-site mutations that suppress the *var2* phenotype has provided substantial support for this hypothesis. It has been concluded that reduced plastidic gene expression can restore the *var2* phenotype. A rational explanation is that these mutations extend the duration of chloroplast development, thereby lowering the threshold level of FtsH activity and suppressing leaf variegation (Yu *et al*., 2004). However, it has also been reported that decreased plastidic gene expression in the suppressor lines can indirectly enhance the level of FtsH activity through an unknown mechanism (Wu *et al*., 2016; Yu *et al*., 2004). Therefore, further investigations are necessary to elucidate the molecular mechanisms underlying the formation of variegated leaves.

Only a few publications have reported that mutants with defective chloroplast division exhibit a leaf variegation phenotype. Kadirjan-Kalbach *et al*. (2012) reported that reduced expression of negative chloroplast division regulator ARC1 encoded by *FtsHi1* led to leaf variegation. In addition, Arabidopsis homologs of the bacterial mechanosensitive (MS) channels of small conductance MscS-Like 2 (MSL2) and MSL3 act as components of the chloroplast division machinery to regulate filamentous temperature-sensitive Z (FtsZ) ring formation (Wilson *et al*., 2011); the *msl2 msl3* double mutant displays a weak leaf variegation phenotype (Haswell and Meyerowitz, 2006). These results indicate that leaf variegation is related to defective chloroplast division. However, the specific role of chloroplast division in variegated-leaf formation has yet to be investigated. We propose that if chloroplast division occurs more slowly than cell division due to genetic mutations or environmental stresses, the number of chloroplasts in daughter cells may diminish, potentially reaching zero. This reduction in chloroplasts could ultimately lead to the development of leaf variegation.

Plastid division is driven by a ring-shaped machinery, which contains the plastid-dividing (PD) ring on the cytosolic side of the outer membrane, the FtsZ ring at the stromal face of the inner membrane, and dynamin-related protein 5B ring (DRP5B ring, also known as ARC5 ring) located at the site of chloroplast division (Osteryoung and Pyke, 2014). The FtsZ ring is first formed by two homologues (FtsZ1 and FtsZ2) of the bacterial division protein FtsZ, which is a self-assembling, microtubulin-like GTPase. The correct placement of the FtsZ ring at the middle of a chloroplast is determined by the Min system composed of MinD1 (minicell D1), MinE1, MCD1 (multiple chloroplast division site1) and ARC3 (Chen *et al*., 2018). The assembly of the FtsZ ring is promoted by ARC6 (Accumulation and Replication of Chloroplasts6), which is localized to the chloroplast inner membrane division site and anchors the FtsZ ring to the site via interaction with FtsZ2, but is inhibited by PARC6 through interaction with ARC3. In addition, the intermembrane space domain of PARC6 and ARC6 recruits the transmembrane proteins PDV1 and PDV2, respectively (Glynn *et al*., 2008; Wang *et al*., 2017; Zhang *et al*., 2016). Finally, DRP5B is recruited by PDV1 and PDV2 and assembled into the DRP5B ring on the cytoplasmic surface of the outer envelope, and constriction initiates (Gao *et al*., 2003; Sun *et al*., 2020). In eukaryotic unicellular algae that own one chloroplast, chloroplast division occurs in the synthesis phase of the cell cycle, and the two duplicated chloroplasts are evenly partitioned into the daughter cells at the mitosis phase (Sumiya *et al*., 2016). In contrast, the relationship between chloroplast division and cell division in higher plants is complex and probably varies according to cell types, developmental stages and environmental conditions (Miyagishima and Kabeya, 2010; Pedroza-Garcia *et al*., 2016). For instance, chloroplasts continuously divide to maintain its stable density during expansion of mesophyll cells (He *et al*., 2021; Miyagishima, 2011), while chloroplasts are partitioned into each daughter cell unbiasedly or stochastically with a tendency toward equality in leaf dividing cells (Birky, 1983; Sheahan *et al*., 2004). Although it is not essential that chloroplasts have to divide before a cell starts division due to several chloroplasts existing in each dividing cell, chloroplast numbers must maintain at a sufficient level during leaf development. However, current knowledge of the regulation of chloroplast biogenesis during leaf development is quite limited.

In this study, we investigated the mechanism underlying leaf variegation formation from the perspective of chloroplast biogenesis. Our results showed that *VAR2* mutations significantly affect both the development and division of chloroplasts during the early stages of leaf development. We found that accelerating either chloroplast division or chloroplast development can suppress leaf variegation. Analyses of chloroplast biogenesis of another leaf variegation mutant *im*, and the chloroplast development mutant *clpR4*, which exhibits a leaf virescent phenotype, further support the notion that chloroplast division plays an important role in variegated leaf formation. Overall, our findings reveal a coordinated yet independent relationship between chloroplast development and division during leaf growth, intricately linking these processes to the phenotype of leaf variegation, virescence, and pale green leaves.

## Results

### *VAR2* mutations result in the majority of cells lacking chloroplasts in the first pair of leaves

To evaluate the effect of *VAR2* mutations on chloroplast biogenesis, we examined the number of chloroplasts during the development of the first pair of leaves. To do this, we made protoplasts from the leaves of 9-, 13- and 17-day-old seedlings grown on half-strength Murashige and Skoog (MS) media containing 1% sucrose under the long day (16 hr light/8 hr dark) condition (Figure 1A). Developmental status of chloroplasts were observed using confocal microscopy via chlorophyll autofluorescence. Our results showed that *var2* protoplasts had more variations in autofluorescence than the wild type (WT) (Figure 1B). To better quantify differential chloroplast development, we classified leaf cells into three categories, type a, b and c, which represent cells with well-developed, poorly developed, and no chloroplasts, respectively, based on chlorophyll autofluorescence (Figure 1C). We found that over 90% of the cells in 9-, 13- or 17-d-old WT plants were classified as type a, whereas only 5%, 12% and 20% of the cells, respectively, were type a in the corresponding *var2* plants (Figure 1C). In addition, the percentage of type c cells in *var2* reached as high as 92% at day 9, whereas no type c cells were detected in WT (Figure 1C). Taken together, our results indicate that *VAR2* mutations severely impair chloroplast biogenesis during leaf development.

**Figure 1.**
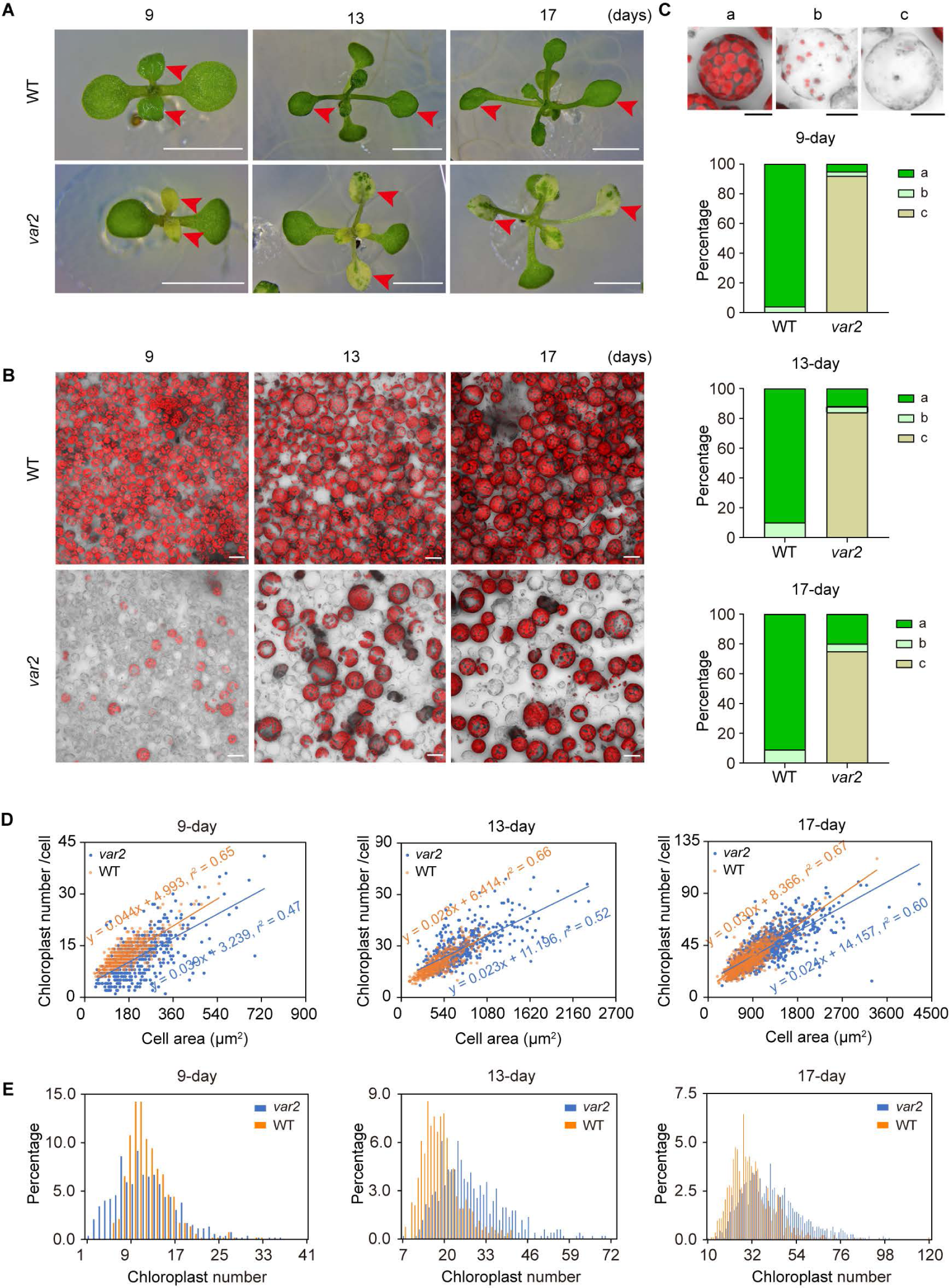
*VAR2* mutations severely inhibit chloroplast biogenesis during leaf development. (A) Phenotypes of 9-, 13-, and 17-day-old WT and *var2* seedlings. Bars, 0.5 cm. Red arrows indicate the first pair of true leaves. (B) Confocal microscopy images of protoplasts isolated from the leaves shown in (A). Red indicates chlorophyll autofluorescence. Bar, 5 μm. (C) Representative protoplasts of type a, b and c, and their corresponding percentage in the total protoplasts. More than 300 protoplasts were randomly selected. Bars, 10 μm. (D) Correlation analysis between chloroplast number and cell size. More than 100 protoplasts with red autofluorescence were analyzed for each genotype. Equations are computed via linear regression. The *r*^2^ (correlation coefficient) value of the best-fit line is shown in each panel. (E) Percentage of cells with the different number of chloroplasts. More than 100 protoplasts were analyzed for each replicate.

### Chloroplast biogenesis is significantly delayed yet prolonged in *var2*

To further investigate the difference in chloroplast biogenesis between WT and *var2*, we examined the correlation between cell size and chloroplast number in type a and b cells. As shown in Figure 1D, the distribution of the chloroplast number per cell over cell size was more dispersed in *var2* than in WT across all three developmental stages, indicating that *VAR2* mutations disturb the uniformity of chloroplasts. Correlation coefficients (*r^2^*) between cell size and chloroplast number per cell were 0.65 and 0.47 in 9-day-old WT and *var2* seedlings, respectively. However these values became closer in 13-day-old (0.66 VS 0.52) or 17-day-old (0.67 VS 0.60) seedlings. In addition, 12.6% of chloroplast-containing cells in *var2*, compared to only 1.7% in WT, were larger than 1500 μm^2^ in the 17-day-old seedlings. These results demonstrate that chloroplast biogenesis is much slower and more heterogeneous in *var2* leaves.

We also analyzed the effect of *VAR2* mutations on the number of chloroplasts per cell over time. In 9-day-old seedlings, 25.3% of the cells contained fewer than 5 chloroplasts in *var2*, compared to only 1.6% in WT (Figure 1E). Conversely, 9.8% of *var2* cells had more than 20 chloroplasts, which was 7% higher than in WT (2.8%) (Figure 1E). A similar trend was observed in 13-day-old seedlings. Interestingly, by 17 days, *var2* cells with an extremely low number of chloroplasts were no longer detected, and the percentage of cells containing more than 50 chloroplasts was significantly higher in *var2* (22.2%) than in WT (10.6%) (Figure 1E). These data indicate that *VAR2* mutations reduce the frequency of plastid division while prolonging the duration of chloroplast biogenesis.

### Defective plastid division leads to the generation of plastid-free cells in *var2*

The above results promoted us to investigate whether plastid division is impaired in *var2*. To address this, we labelled plastids by expressing plastid ribosomal protein L11 (PRPL11)-GFP fusion protein in *var2* (*PRPL11*-*GFP*/*var2*), which exhibits similar phenotypes of leaf variegation and comparable percentages of three-type cells as *var2* (Figure S1). We analyzed plastid numbers in individual protoplasts isolated from the green and yellow sectors of *PRPL11*-*GFP*/*var2* mature variegated leaves. In the green sector, 86.83% of the cells contained chloroplasts, showing both chlorophyll autofluorescence and GFP fluorescence, while 10.48% of the cells displayed only GFP signals, indicating the presence of plastids without chlorophyll, such as leucoplasts. In contrast, only 21.05% of the cells in the yellow sector owned chloroplasts, while 62.87% contained only GFP signals (Figure 2A, 2B). Surprisingly, neither GFP nor red chlorophyll autofluorescence were observed in some cells, suggesting the presence of plastid-free cells. The percentages of plastid-free cells were 14.04% in the yellow sector and 0.90% in the green sector (Figure 2B).

**Figure 2.**
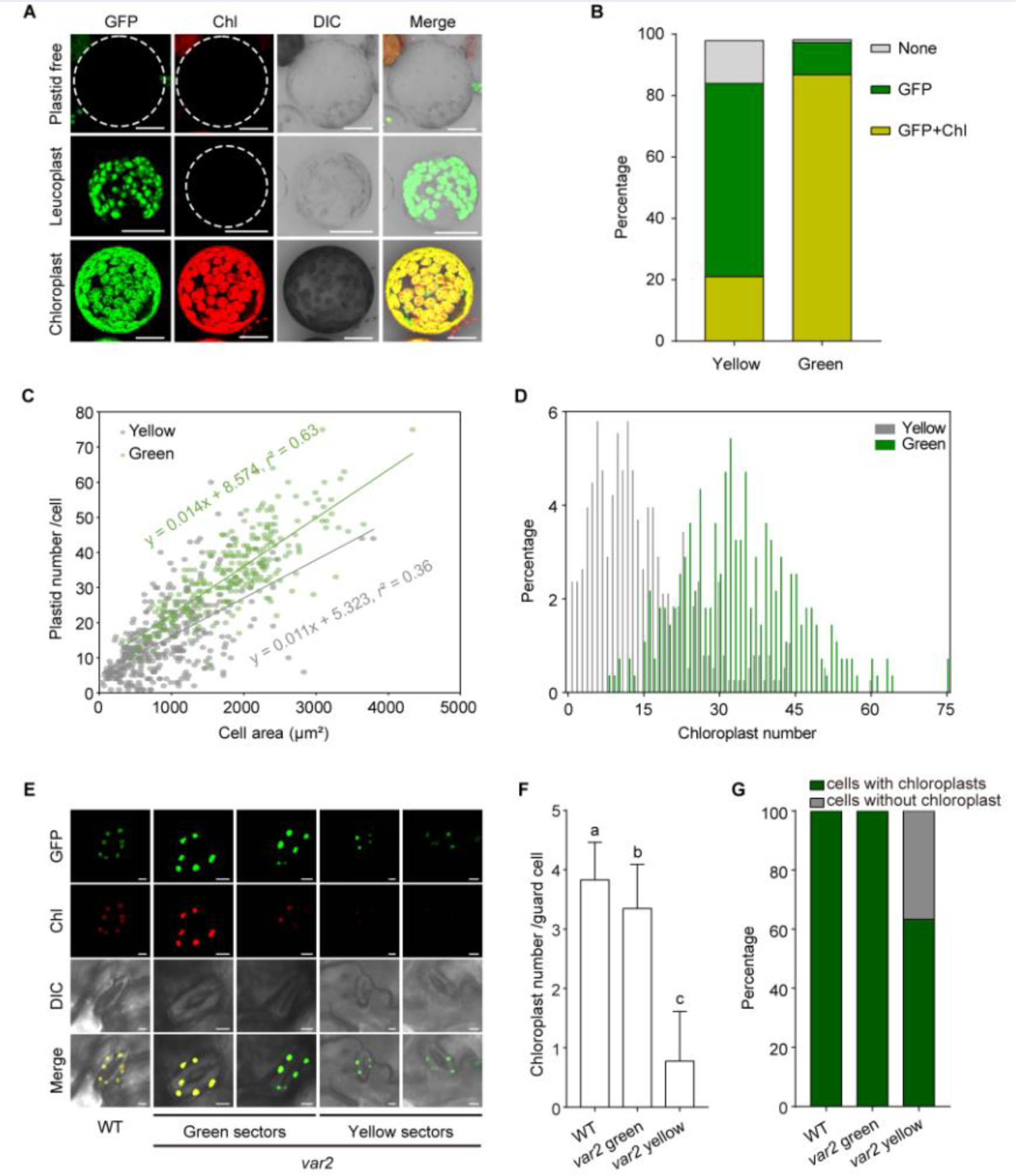
*VAR2* mutations impair chloroplast division and result in the formation of plastid-free cells. (A) Confocal microscopy analysis of protoplasts isolated from green and yellow sectors of mature leaves of *PRPL11*-*GFP*/*var2* plants grown in soil. White circles indicate plastid-free cells that lack both red autofluorescence and GFP signals. GFP, green fluorescence protein; Chl, chlorophyll autofluorescence; DIC, differential interference contrast microscopy. Bars, 20 μm. (B) Percentage of the three types of cells with chloroplasts, leucoplasts or without plastids shown in (A). More than 300 protoplasts were analyzed. (C) Correlation analysis between chloroplast number and cell size in green and yellow sectors of *var2*. More than 150 protoplasts were analyzed per replicate. The *r*^2^ values of the best-fit lines are 0.63 (green sectors) and 0.36 (yellow sectors). (D) Distribution of cells containing different chloroplast number in green and yellow sectors of *var2*. More than 150 protoplasts were analyzed per replicate. (E) Confocal microscopy images of guard cells in *RPL11-GFP/*WT and *RPL11-GFP/var2* leaves. Bars, 4 μm. (F) Average of chloroplast number per guard cell. Significant differences between genotypes were marked with different letters (One-Way ANOVA, *P* < 0.05). (G) Percentage of cells with or without chlorophyll autofluorescence. More than 150 guard cells were analyzed in (F) and (G).

We further quantified the number of chloroplasts and size per protoplast isolated from green and yellow sectors of *var2* mature leaves. The average cell size in the yellow sector was 893.5 μm^2^, significantly smaller than 1846.1 μm^2^ observed in the green sector. Consistently, the chloroplast number per cell was markedly lower in the yellow sector. Notably, 75.7% of the cells in the yellow sector possessed 1 to 20 chloroplasts, compared to only 11.6% in the green sector (Figure 2C, 2D). Overall, our findings reveal that *VAR2* mutations impair plastid division, ultimately leading to the formation of plastid-free cells.

### Chloroplast division is also inhibited in *var2* guard cells

Since chloroplast numbers can be easily calculated in guard cells, we investigated whether *VAR2* mutations have an impact on chloroplast division in guard cells. To do this, we introduced *PRPL11*-*GFP* into the WT genetic background by crossing with *PRPL11*-*GFP*/*var2*. Our data showed that introduction of *PRPL11-GFP* into WT had no significant effect on chloroplast biogenesis (Figure S1). Chloroplast numbers in guard cells were analyzed in mature leaves of 20-day-old *PRPL11*-*GFP*/WT and *PRPL11*-*GFP*/*var2* plants grown in soil. The average number of chloroplasts per guard cell were 3.35 in *var2* green sectors, which were significantly less than in WT (3.83), and only 0.75 in the yellow sector (Figure 2E, 2F). In addition, all guard cells in the green sector contained chloroplasts, while only 63.4% of guard cells in the yellow sector had chloroplasts (Figure 2G). Interestingly, plastid-free guard cells were not detected in *var2*, unlike in leaf cells,. Taken together, our data demonstrate that *VAR2* mutations impair chloroplast division in guard cells as well.

### Cell division is not affected in *var2*

Although our data demonstrated that some *var2* cells are devoid of plastids due to defective chloroplast biogenesis during leaf development, we cannot exclude the possibility that *VAR2* mutations also affect cell division. To investigate this, we measured cell numbers in the first pair of true leaves and root meristems. As shown in Figure 3A and 3B, no significant differences in cell number were observed between the leaves of 17-day-old WT and *var2* plants. Similarly, no obvious differences in root meristem size or meristem cell number were detected between 7-day-old WT and *var2* seedlings (Figure 3C-3E). These results indicate that cell division occurs normally in *var2*.

**Figure 3.**
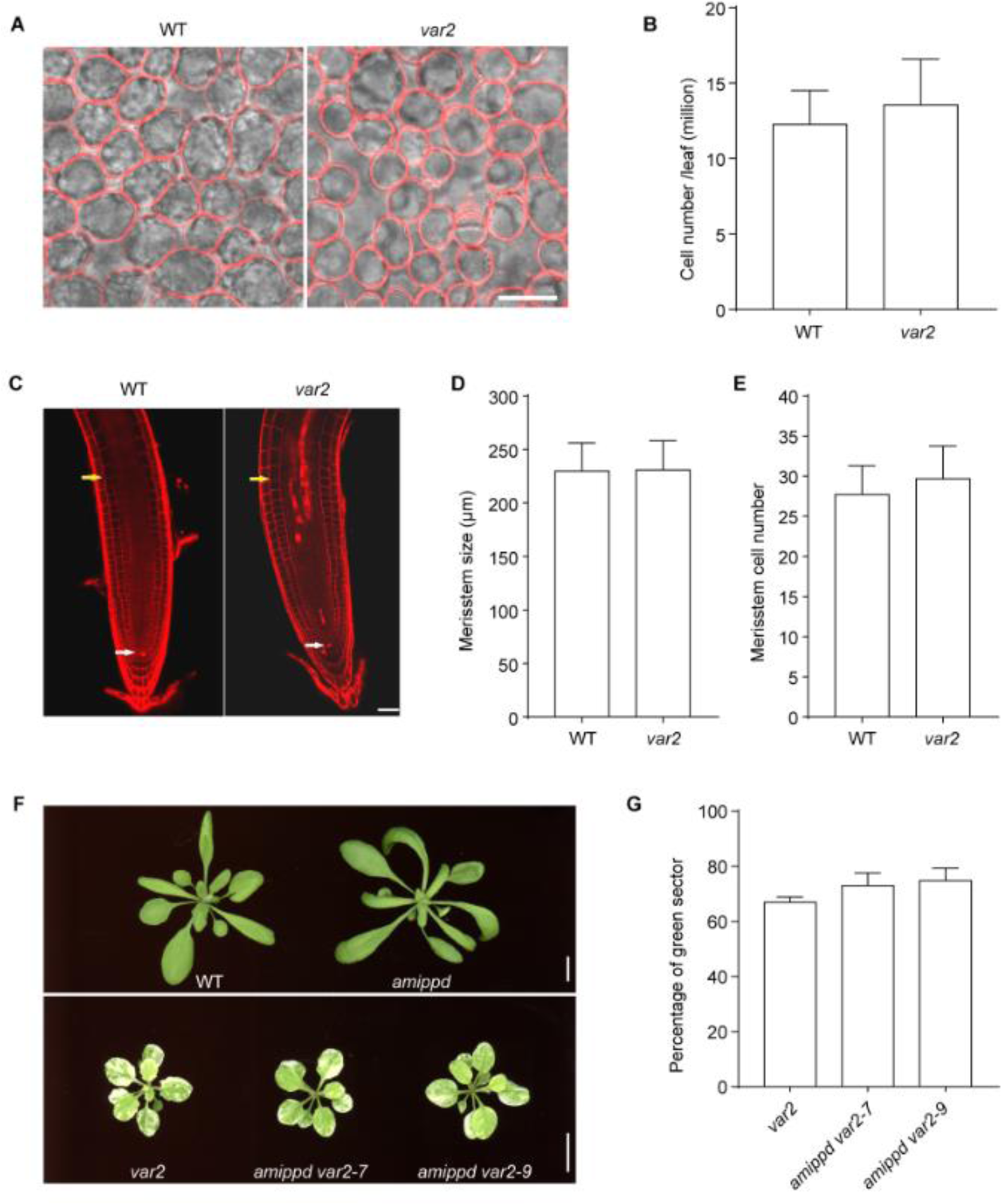
Cell division is not affected by *VAR2* mutation. (A) Confocal microscopy images of WT and *var2* leaf cells stained with FM4-64. Bar, 40 μm. (B) Mesophyll cell number in WT and *var2* leaves. Data were means ± SD (n = 30). (C) Confocal microscopy images of WT and *var2* root tips stained with propidium iodide. White and yellow arrows indicate quiescent centers and the ends of meristem zone, respectively. Bar, 20 μm. (D) and (E) are the size and cell number of root meristems. Data were means ± SD (n = 40). (F) Phenotypes of 20-day-old WT, *amippd*, *var2*, and two independent lines of *amippd var2* plants grown in soil. Bars, 1 cm. (G) Percentage of green sectors in *var2* and *amippd var2* plants. Data were means ± SD (n = 3).

To further explore whether accelerated cell proliferation affects the leaf variegation phenotype of *var2*, we knocked down the expression of *Peapod* (*PPD*), which suppresses the proliferation of dispersed meristematic cells in leaves, in the *var2* background (*amippd*/*var2*) using the artificial microRNA technique (Schwab *et al*., 2006). Surprisingly, *amippd/var2* displayed a weaker leaf variegation phenotype and a higher percentage of green sectors, but no significant difference in the phenotype was observed compared to *var2* (Figure 3F, 3G). Taken together, our data indicate that cell division is not affected by *VAR2* mutations.

### Chloroplast division is involved in the formation of leaf variegation

To investigate the role of chloroplast biogenesis in variegated leaf formation, we first examined whether plastid division contributes to leaf variegation of *var2*. We analyzed the expression levels of key plastid-division genes, such as *ARC6*, *PARC6*, *PDV1* and *PDV2*, in *var2* leaves during different stages of development. Quantitative PCR (qPCR) analysis showed that expression levels of these genes were not significantly different between WT and *var2* at early stages of leaf development but appeared to decrease more rapidly in *var2* than WT at later stages (Figure S2), indicating that *VAR2* mutations have no substantial effect on mRNA levels of plastid division genes, particularly during early leaf development.

To provide genetic evidence for the involvement of chloroplast division in leaf variegation, we made double mutants by crossing *var2* with *arc6*, *parc6*, *pdv1* or *pdv2* mutants (Figure S3). Phenotypic analysis showed that the single mutants (*arc6*, *parc6*, *pdv1* and *pdv2*) displayed green leaves similar to WT (Figure 4A, 4B). Among the double mutants, *parc6 var2* and *pdv1 var2* displayed much more severe leaf variegation compared to *var2*, whereas *arc6 var2* and *pdv2 var2* had similar variegation to *var2* at both the seedling (Figure 4A) or juvenile stages (Figure 4B). Quantification analysis confirmed that the percentage of the green area in *parc6 var2* and *pdv1 var2* leaves was significantly reduced, compared to *var2,* while no difference was observed between *var2* and *arc6 var2* or *pdv2 var2* had (Figure 4C and 4D). These result suggest that *PARC6* and *PDV1* play critical roles in the formation of leaf variegation in *var2*.

**Figure 4.**
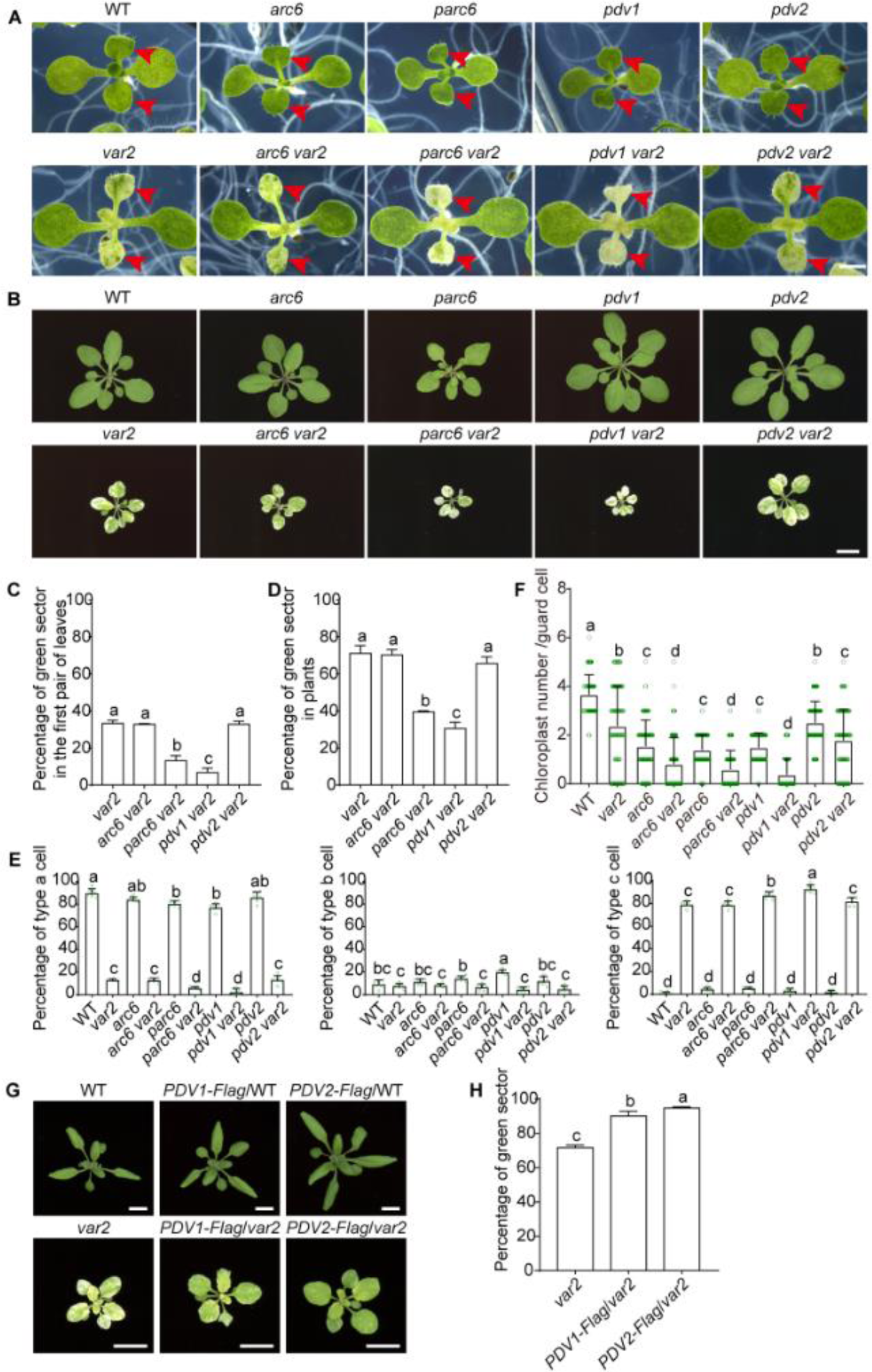
Chloroplast division is involved in the formation of leaf variegation. (A) and (B) Phenotypes of 13-day-old (A) and 20-day-old (B) WT, *var2*, *arc6*, *parc6*, *pdv1*, *pdv2*, *arc6 var2*, *parc6 var2*, *pdv1 var2*, and *pdv2 var2* plants. Bar in (A), 1 mm. Bar in (B), 1 cm. Red arrows in (A) indicate the first pair of true leaves. (C) and (D) Percentage of green sectors in the first pair of leaves shown in (A) and in plants shown in (B), respectively. Data were means ± SD (n = 3). (E) Percentage of type a, b and c cells isolated from the first pair of leaves shown in (A). Data were means ± SD (n = 6). (F) Chloroplast number per guard cell in green sectors of plants shown in (B). A total of 100 guard cells were analyzed. (G) Overexpression of *PDV1* and *PDV2* significantly rescues the *var2* phenotype. PDV1 or PDV2 fused with Flag was overexpressed in WT and *var2* genetic backgrounds. The shown were 20-day-old plants grown in soil. Bars, 1 cm. (H) Percentage of green sectors in plants shown in (G). Data were means ± SD (n = 3). Statistical analysis in all experiments was performed with One-Way ANOVA, and significant differences among genotypes were marked with different letters (*P* < 0.05).

To further evaluate the effects of mutations in *PARC6*, *ARC6, PDV1* and *PDV2* on chloroplast biogenesis, we analyzed protoplasts isolated from the first pair of leaves. Confocal microscope analysis showed that the proportion of type a cells, which contain well-developed chloroplasts, was significantly lower in *parc6 var2* and *pdv1 var2* double mutants than in *var2*, while it remained unchanged in *arc6 var2* and *pdv2 var2* (Figure 4E). In contrast, the percentage of type c cells, which lack chloroplasts, was markedly higher in *parc6 var2* and *pdv1 var2* than in *var2*, whereas it was comparable to *var2* in *arc6 var2* and *pdv2 var2* (Figure 4E). No significant difference in the proportion of type b cells was detected between *var2* and any of the double mutants (Figure 4E). In addition, the number of chloroplasts in guard cells of all double mutants were significantly reduced, compared to *var2* (Figure 4F). Taken together, these findings confirm that chloroplast division is impaired in *var2*, and mutations in *PARC6* and *PDV1* exacerbate the variegated phenotype of *var2*.

Simultaneously, we tested whether accelerating chloroplast division could improve the leaf variegation phenotype of *var2*. Since previous studies have shown that overexpression of *PDV1* and *PDV2* increases chloroplast numbers in mesophyll cells (Miyagishima *et al*., 2006), we generated transgenic plants overexpressing either *PDV1* or *PDV2* in the *var2* background (Figure S4). As expected, transgenic plants overexpressing *PDV1* (*PDV1*-*Flag*/*var2*) or *PDV2* (*PDV2*-*Flag*/*var2*) significantly mitigated the variegated leaf phenotype of *var2* (Figure 4G). The area of green sectors in *PDV1*-*Flag*/*var2* and *PDV2*-*Flag*/*var* plants was approximately 90% and 95%, respectively, which was significantly greater than the 71.68% observed in *var2* plants (Figure 4H). Notably, while *PDV2* mutation did not affect leaf variegation in *var2*, overexpression of *PDV2* markedly improved the *var2* phenotype, suggesting that *PDV1* (a homolog of *PDV2*) may play a more critical role in plastid division in *var2*. Collectively, our data indicate that chloroplast division is involved in leaf variegation of *var2*.

### Early chloroplast biogenesis is critical for suppression of leaf variegation

Genetic screening for suppressors of leaf variegation has identified many suppressor genes, such as *Plastid Ribosomal Protein L11* (*PRPL11*) and *Suppressor of Thylakoid Formation1* (*SOT1*), which are known to regulate chloroplast gene expression (Wu *et al*., 2016; Wu *et al*., 2013). To inspect the role of the suppressor genes in chloroplast biogenesis, we analyzed the relationship between chloroplast number and cell size in the first pair of leaves from 13-day-old WT, *var2*, *prpl11* and *prpl11 var2* seedlings (Figure 5A). Our results showed that type a cells with well-developed chloroplasts accounted 90.5% of cells in *prpl11*, a proportion comparable to that of WT (92.6 %), whereas *prpl11 var2* had a significantly higher proportion of type a cells (76.2%) than *var2* (12.4 %). Notably, type c cells lacking chloroplasts were absent in *prpl11 var2* (Figure 5B). These results indicate that *PRPL11* mutations substanti ally improve chloroplast biogenesis in *var2*. When chloroplast number was plotted against cell size, the correlation coefficient (*r^2^*) for *prpl11 var2* (0.74) and *prpl11* (0.77) were significantly higher than that of *var2* (0.52) (Figure 5C). Interestingly, both *prpl11* and *prpl11 var2* exhibited a steeper slope in the regression equation than WT, indicating that cells of the same size in these genotypes contain more chloroplasts than WT. Similar trends were observed in another suppressor of leaf variegation, *sot1* (Figure S5). Thus, our data underscore the importance of early chloroplast biogenesis in mitigating the variegation phenotype.

**Figure 5.**
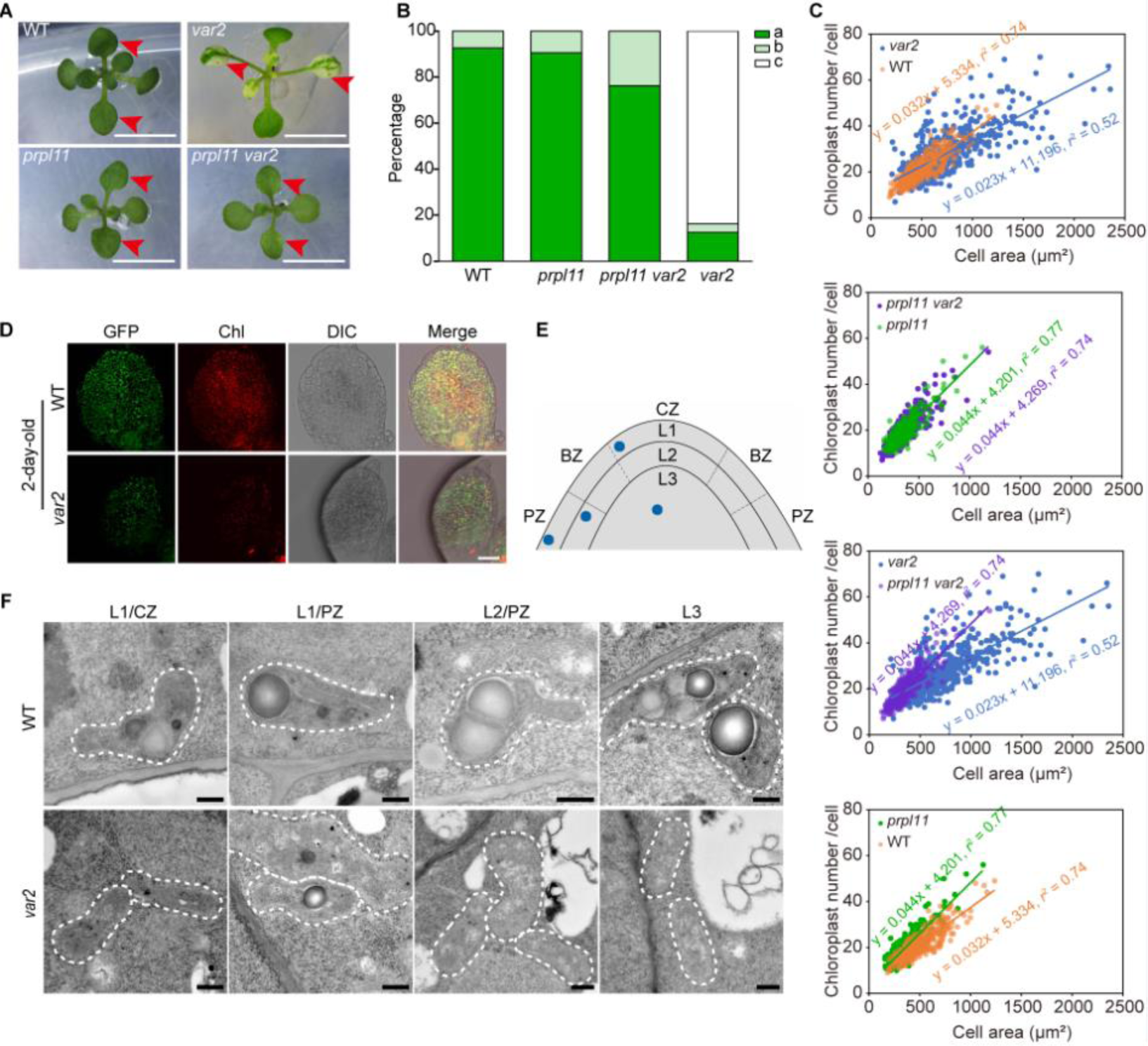
Early chloroplast biogenesis is critical for suppression of leaf variegation. (A) Phenotypes of 13-day-old WT, *var2*, *prpl11* and *prpl11 var2* seedlings. Red arrows indicate the first pair of true leaves. Bars, 0.5 cm. Red arrows indicate the first pair of leaves. (B) Percentage of type a, b, or c cells isolated from the first pair of leaves. More than 400 protoplasts were analyzed in each sample. (C) Correlation analysis between chloroplast number and cell area in type a cells. More than 350 cells were analyzed for each sample. Correlation coefficients (*r*^2^) are shown in the panel. (D) Confocal microscopy images of the first pair of leaves from 2-day-old *PRPL11-GFP*/WT and *PRPL11-GFP*/*var2* seedlings. Green signals indicate plastids; red signals indicate chlorophyll autofluorescence. Bar, 30 μm. (E) Schematic drawing of the SAM. Blue dots represent positions of tomographic pictures shown in (F). CZ, the central zone; BZ, the zone between the CZ and PZ; PZ, the peripheral zone. L1, L2 and L3 indicate the three layers of the SAM. (F) TEM analysis of the transition from proplastids into chloroplasts in the SAMs of 3-day-old WT and *var2* seedlings cultured on half-strength MS media with 1% sucrose. Bars, 500 nm.

To validate the role of *VAR2* in early chloroplast biogenesis, we examined chloroplast numbers and development in the first pair of leaves using transgenic plants expressing PRPL11–GFP in WT and *var2*. Our results showed that in *var2* both chlorophyll autofluorescence intensity and GFP signal density were markedly reduced in whole leaves from 2-day-old seedling, compared to WT (Figure 5D). Moreover, a significant proportion of cells in *var2* leaves contained only GFP signals (Figure 5D), indicating an increased prevalence of the non-photosynthetic plastids. Thus, these results confirm that chloroplast biogenesis at the early stage of leaf development is severely inhibited by *VAR2* mutations.

### Proplastid-to-chloroplast transition is inhibited in the SAM of *var2*

It has been demonstrated that most proplastids in the shoot apical meristem (SAM), particularly in the L1 and L3 layers, are partially differentiated into chloroplasts (Charuvi *et al*., 2012). To determine whether *VAR2* mutation affects the transition of proplastids to chloroplasts in the SAM, we analyzed plastid ultrastructure using transmission electron microscopy (TEM). In 3-day-old *var2* seedlings grown on half-strength MS media with 1% sucrose in the 16 hr light/8 hr dark cycle, TEM analysis showed that plastids in *var2* cells had fewer thylakoid networks compared to those in WT cells occupying equivalent SAM positions (Figure 5E, 5F). In the central zone (CZ), plastids in both L1 and L3 layers of WT cells exhibited distinct thylakoid membranes, whereas plastids in *var2* cell displayed a near-complete absence of these structures (Figure 5F). In the peripheral zone (PZ), WT plastids from the L1 and L2 layers contained well-developed thylakoid membranes, whereas *var2* plastids from the same layers were characterized by the presence of vesicles and tubules (Figure 5F). These results clearly indicate that *VAR2* mutations significantly inhibit chloroplast development at a very early stage, affecting the transition from proplastids to chloroplasts within the SAM.

### Pchlide synthesis is defective in etiolated seedlings of *var2*

Chlorophyll synthesis is a crucial prerequisite for chloroplast development. It remains unknown if *VAR2* mutations affect chlorophyll biosynthesis. To solve this question, we examined the greening process in etiolated *var2* seedlings. Interestingly, 4-day-old *var2* etiolated seedlings displayed a pale green coloration after exposure to light, in contrast to the greening cotyledon observed in WT seedlings (Figure 6A). Chlorophyll content in *var2* seedlings was significantly lower than in WT after 30 min of de-etiolation (Figure 6B). Given the compromised chlorophyll synthesis, we postulated that the synthesis of protochlorophyllide (Pchlide), a precursor of chlorophyll, was blocked in *var2* etiolated seedlings. As expected, Pchlide levels in 4-day-old *var2* etiolated seedlings were approximately 50% of those in WT (Figure 6C). Furthermore, Pchlide content in *var2* seedlings remained relatively stable following 30-min of light exposure, whereas it rapidly declined in WT seedlings (Figure 6C). These findings suggest that VAR2 is essential for Pchlide synthesis in the dark.

**Figure 6.**
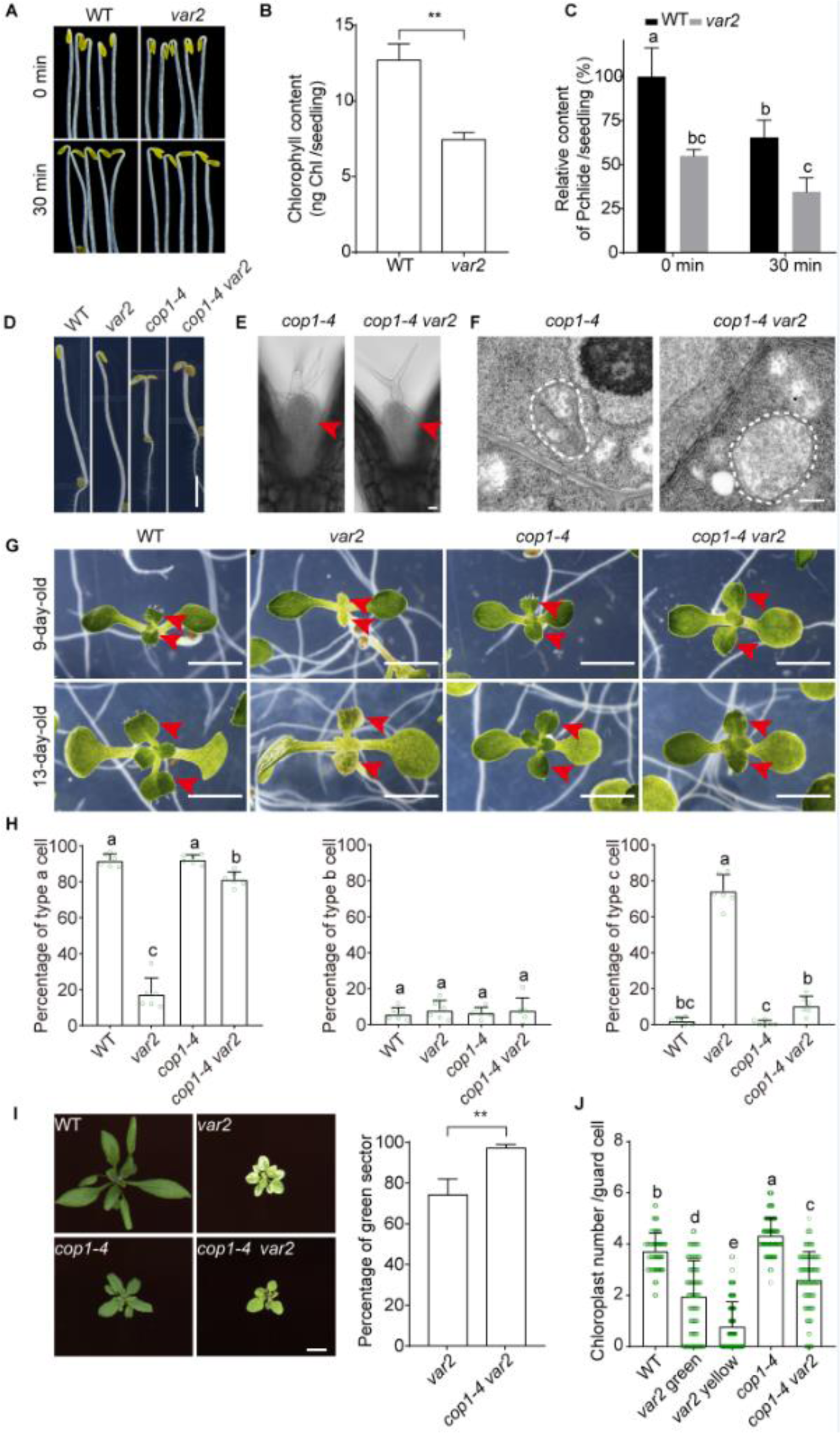
*COP1* mutations fully rescue leaf variegation in *var2*. (A) Phenotypes of 4-day-old WT and *var2* etiolated and de-etiolated (for 30-min) seedlings. Bar, 1 mm. (B) Chlorophyll content of 4-day-old etiolated seedlings exposed to light for 30 min. The data were means ± SD (n = 3). Significant difference between WT and *var2* was labelled with stars (student *t-*tes, *P* < 0.01). (C) Relative Pchlide content per seedling shown in (A). The data were means ± SD (n = 3). (D) Phenotypes of 4-day-old WT, *var2*, *cop1-4* and *cop1-4 var2* etiolated seedlings. Bar, 1 mm. (E) Confocal microscopy images of the first pair of leaves (labelled with red arrows) from *cop1-4* and *cop1-4 var2* etiolated seedlings. Bar, 30 μm. (F) TEM images of plastids in the first pair of leaves shown in (E). Bar, 500 nm. (G) Phenotypes of 9- and 13-day-old WT, *var2*, *cop1-4* and *cop1-4 var2* seedlings. Bars, 1 mm. Red arrows indicate the first pairs of leaves. (H) Percentage of type a, b and c cells from the first pair of leaves from 13-d-old seedlings shown in (G). The Data were means ± SD (n = 6). (I) Phenotypes of 20-day-old WT, *var2*, *cop1-4*, and *cop1-4 var2* plants grown in soil in the 16 hr light/8 hr dark cycle, and the percentage of green sectors in *var2* and *cop1-4 var2* plants. Bar, 1 cm. The data were means ± SD (n = 3). Stars indicate significant difference between *var2* and *cop1-4 var2* (student *t-*tes, *P* < 0.01). (J) Chloroplast number per guard cell in plants shown in (I). A total of 130 guard cells were analyzed. Statistical analyses in (C), (H) and (J) were performed with One-Way ANOVA, and significant differences among genotypes were marked with different letters (*P* < 0.05).

### *COP1* mutations fully rescue leaf variegation of *var2*

COP1 is a central repressor of photomorphogenesis, and its mutations lead to photomorphogenic development, such as open and expanded cotyledons, short hypocotyls, and partially developed thylakoids in darkness (Han *et al*., 2020). To examine whether *COP1* mutations affect chloroplast biogenesis in the *var2* background, we generated the *cop1-4 var2* double mutant through genetic crossing. Our data showed that 4-day-old etiolated seedlings of *cop1-4 var2* displayed a constitutive photomorphogenic phenotype similar to those of *cop1-4*, characterized by open cotyledons and shorter hypocotyls. In contrast, *var2* and WT etiolated seedlings exhibited a typical skotomorphogenic phenotype with closed cotyledons and elongated hypocotyls (Figure 6D). In addition, the first pair of leaves emerged in 4-day-old *cop1-4* and *cop1-4 var2*, but not in *var2* and WT etiolated seedlings (Figure 6E). TEM analysis showed partially developed thylakoid structures in leaves of both *cop1-4* and *cop1-4 var2* in darkness (Figure 6F). Under normal light conditions, *cop1-4 var2* seedlings exhibited almost no leaf variegation at 9 and 13 days old when grown on half-strength MS media containing 1% sucrose (Figure 6G). Consistently, microscope analysis of protoplasts from the first pair of leaves of 13-day-old seedlings revealed significant changes in the percentages of type a and c cells in *cop1-4 var2*. Type a cells with well-developed chloroplasts were significantly higher in *cop1-4 var2* than in *var2*, while type c cells were markedly reduced, compared to *var2* (Figure 6H). Type B cells remained unaffected by the *COP1* mutation in the *var2* background (Figure 6H). When grown in soil, *cop1-4 var2* plants exhibited almost no leaf variegation phenotype (Figure 6I). The area of green sectors in leaves was significantly higher in *cop1-4 var2* than in *var2* (Figure 6I). In addition, the number of chloroplasts in *var2* guard cells was also significantly rescued by *cop1-4* (Figure 6J). Finally, we investigated whether *COP1* mutations could rescue the variegated leaf phenotype of *var2* under various photoperiods. Across all tested light cycle conditions, *cop1-4* almost fully suppressed the *var2* phenotype (Figure S6). Taken together, our data suggest that *COP1* mutations facilitate chloroplast biogenesis and rescue the leaf variegation phenotype of *var2* under both dark and light conditions.

### Impaired chloroplast division as a common feature in variegation mutants but not in virescent mutants

To determine whether defective plastid division is a common feature of other variegated mutants, we investigated chloroplast biogenesis in *im*, which encodes a chloroplast terminal oxidase. The *im* mutant exhibited more severe leaf variegation than *var2*, as demonstrated by a significant reduction in the area of green sectors (Figure 8A, 8B). Like in *var2*, the average number of chloroplasts per guard cell was significantly lower in both the green sector and yellow sector of *im*, compared to WT (Figure 8C). These results suggest that plastid division is impaired in *im*, as in var2.

Mutants with defective chloroplast biogenesis can display a range of leaf coloration phenotypes, such as variegation, virescence, and pale green. However, it remains unclear whether virescent mutants are defective in plastid division. To address this, we examined chloroplast development and numbers in the virescent mutant *clpr4* (Figure 8D-8G). Microscopy analysis of isolated protoplasts from the first pair of leaves showed that *clpr4* exhibited a distribution of cell types similar to WT, predominantly consisting of type a cells. This pattern contrasts sharply with *var2*, which showed a majority of type c cells devoid of chloroplasts (Figure 8E). To investigate whether *ClpR4* mutations affect plastid division, we quantified the number of chloroplasts in guard cells. Interestingly, *clpr4* had the significantly higher number of chloroplasts than WT and *var2* (Figure 8G). These results suggest that enhanced chloroplast division in *clpr4* compensates for its defects in chloroplast development.

## Discussion

Over the past decades, studies on VAR2-mediated leaf variegation have primarily focused on elucidating the regulatory network of chloroplast development via genetic screening and functional analysis of suppressor and enhancer genes. However, the impact of *VAR2* mutations on chloroplast division has remained largely overlooked. Here, we found dual roles of VAR2 in both chloroplast development and division, providing new insights into the mechanisms underlying variegated leaf formation in *Arabidopsis thaliana*.

### VAR2 is essential for both development and division of chloroplasts

*VAR2* mutations severely impair chloroplast biogenesis during early leaf development, leading to delayed chloroplast maturation and an increased proportion of cells with non-photosynthetic plastids. In addition, the biogenesis process is prolonged in *var2*, resulting in greater variability in chloroplast number per cell and cell size. VAR2 also plays a pivotal role in the proplastid-to-chloroplast transition in the SAM, where its absence significantly disrupts thylakoid network formation. Thus, these findings underscore the fundamental role of VAR2 in establishing chloroplasts during early leaf development.

Contrary to previous reports that indicated that the green sector of *var2* are comparable to those in WT (Sakamoto, 2003), we found that the green area in *var2* harbor more and larger chloroplasts than that in WT, suggesting that green sectors may produce more photosynthate to meet the metabolic demands of the yellow sectors. In contrast, non-photosynthetic cells dominate the yellow sector due to defects in chloroplast development (Kato *et al*., 2007). Consistently, our quantitative analysis showed that about 80% of cells in the yellow sectors are non-autotrophic (Figure 2). It is proposed that misfolded and photodamaged proteins will accumulate in the absence of *VAR2*, and thylakoid membrane formation will be blocked at the early stage of chloroplast development, ultimately resulting in chloroplast dysfunction (Dogra *et al*., 2019). However, investigation has intensively focused on the role of *VAR2* in maintaining homeostasis of PSII activity (Kato *et al*., 2018; Kato *et al*., 2023). Therefore, it remains unclear how *VAR2* mutations influence chloroplast development. In this study, we discovered that *VAR2* mutations suppress de-etiolation of cotyledons due to less accumulation of Pchlide and its slower conversion in light, suggesting that VAR2 has distinct function in the dark from that in light. These findings provide a new approach to explore molecular mechanism by which VAR2 regulates chloroplast development.

The L1 and L2 layers of the SAM give rise to the epidermis and outer mesophyll of a leaf, respectively, while the L3 layer contributes to the inner mesophyll and vasculature (Tsukaya, 2002). The transition of proplastids into chloroplasts initiates across these layers, but thylakoid membranes are small, partially differentiated in the L1, L3 and PZ cells (Charuvi *et al*., 2012; Yadav *et al*., 2019). These thylakoid networks keep till the young leaf primordium. Typical thylakoid ultrastructure with granal and stromal membranes appears in the old leaf primordium, where heterogeneity of chloroplast development is obvious due to different cell types (Charuvi *et al*., 2012). Our TEM analysis showed that thylakoid network formation was significantly inhibited in the SAM of *var2*, leaving most plastids in a proplastids state (Figure 5F). This generally aligns with previous findings (Sakamoto *et al*., 2009) but challenges the assertion that differential thylakoid networks in WT and *var2* PZs have no impact on chloroplast development in subsequent leaf development. Since the leaf primordium emerges from the PZ, we hypothesize that *VAR2* mutations have an impact on chloroplast development in the young leaf primordia. This hypothesis is consistent with our genetic evidence showing that accelerating chloroplast biogenesis rescues the variegated phenotype of *var2* (Figure 4G, 6I), highlighting the importance of timely chloroplast development in mitigating leaf variegation.

Following the initiation of leaf primordia, the establishment of leaf polarities (dorsoventral, proximodistal and mediolateral) and subsequent development into the flattened leaf blade and petiole occur, accompanied by rapid cell division (Tsukaya, 2013). Thus, it is important to keep chloroplast biogenesis in pace with cell division at early leaf development. Our results revealed that the yellow sector of *var2* leaves predominantly consist of non-photosynthetic cells, which either lack chloroplasts or any kinds of plastids.

Generally, the active degeneration of thylakoids is restricted to the maturation of leaf pavement cells and does not occur during the development of young leaf cells (Charuvi *et al*., 2012). A plausible explanation for the absence of chloroplasts in *var2* cells is a self-protection mechanism aimed at mitigating photodamage caused by reactive oxygen species (ROS) generated from photosynthesis. This process likely leads to the formation of non-photosynthetic plastids and is in agreement with the observations that variegation severity is proportional to light intensity (Rosso *et al*., 2009). In addition, approximately 15% of cells in the yellow sector lack plastids entirely (Figure 2B). This phenomenon may stem from two potential mechanisms: (1) the rate of chloroplast division may lag behind that of cell division, resulting in plastid-free cells; or (2) damaged chloroplasts might be actively removed by autophagy pathways (Niwa et al., 2004; Izumi et al., 2017). Together, these mechanisms provide a plausible explanation for the development of the yellow sector in *var2* leaves.

### An important role of chloroplast division in variegated leaf formation

One possible explanation for why most chloroplast division-related mutants do not exhibit variegation or virescent leaf phenotypes is that developmentally normal chloroplasts in these mutants can propagate through budding rather than binary fission (Forth and Pyke, 2006; Robertson *et al*., 1995). In *var2*, however, mutations disrupt both chloroplast development and division. Genetic interactions with division-related genes, such as *PDV1* and *PARC6*, highlight the importance of chloroplast division in the formation of variegated leaves in *var2*. For instance, mutations in these genes exacerbated the variegation phenotype, while overexpression of *PDV1* or *PDV2* alleviated it. Interestingly, despite these defects, proplastid division in the SAM and RAM of *var2* remains unaffected, suggesting that VAR2 does not participate in synchronizing proplastid division with the cell cycle. Large plastid nucleoids observed in the yellow sectors of various plant species (Sakamoto *et al*., 2009) might result from defective plastid division, similar to cellular DNA endoduplication. Kato *et al*. (2007) reported that all cells in the yellow sectors of *var2* retain undifferentiated plastids, which contradicts our findings. We observed that approximately 15% of cells isolated from the yellow sectors of mature *var2* leaves lacked any plastids (Figure 2B). This discrepancy may arise from differences in methods and materials. While Kato *et al*. (2007) examined fully expanded leaves from 7- or 8-week-old plants, we analyzed protoplasts from the yellow sectors of the first pair of leaves, which exhibit the most pronounced variegation. Our protoplast-based approach may have increased the likelihood of detecting abnormal cells in *var2*. The presence of plastid-devoid cells has also been reported in *crl* mutants, which display pale green and variegated leaf phenotypes due to insufficient chloroplast division (Asano *et al*., 2004; Chen *et al*., 2009). Furthermore, consistent with reports that aplastidic cells cannot complete the cell cycle (Hudik *et al*., 2014), we found that the proportion of plastid-lacking cells in the whole leaf was minor (Figure 2B).

Chloroplast division defects are also evident in other variegated mutants, such as *im*. In the yellow sectors of *im*, guard cells contained significantly fewer chloroplasts compared to *var2* (Figure 7C), suggesting that impaired chloroplast division is a common feature of variegated mutants. Conversely, the virescent mutant *clpr4*, despite its defective chloroplast development, exhibits accelerated chloroplast division compared to WT (Figure 7G). This increased division rate ultimately restores leaf color to WT levels, indicating that enhanced chloroplast division can partially compensate for reduced chloroplast biogenesis caused by developmental defects. Collectively, our findings strongly suggest that defective chloroplast division plays a crucial role in the formation of variegated leaves.

**Figure 7.**
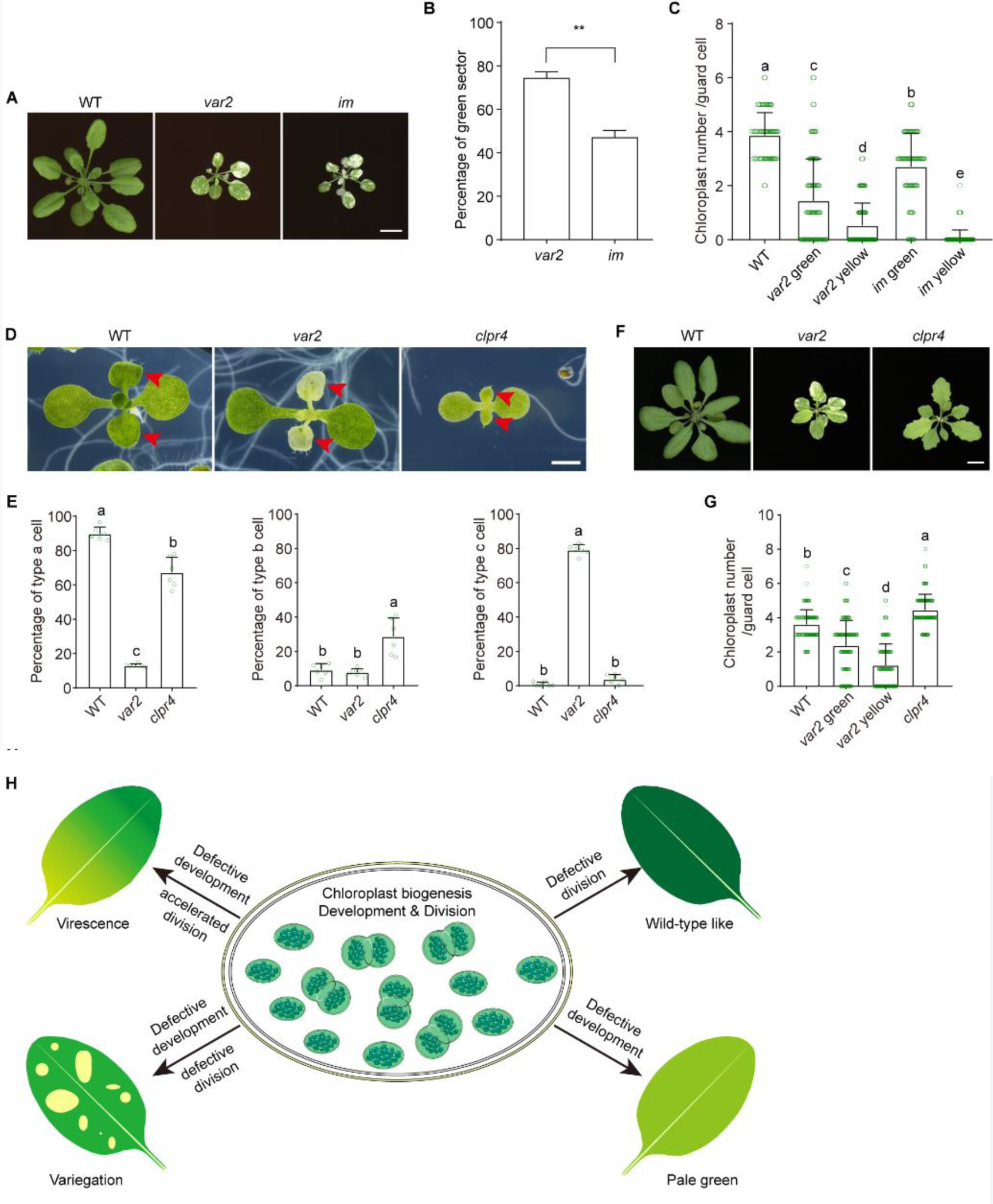
Chloroplast division is a key factor determining leaf variegation. (A) Phenotypes of 20-day-old WT, *var2* and *im* plants grown in soil. Bar, 1 cm. (B) The percentage of green sectors in leaves shown in (A). Data were means ± SD (n = 3). Stars indicate significant difference between *var2* and *im* (student *t*-tes, *P* < 0.01). (C) Chloroplast number per guard cell in plants shown in (A). A total of 100 guard cells were analyzed. Significant differences among genotypes were marked with different letters (One-Way ANOVA, *P* < 0.05). (D) Phenotypes of 13-day-old WT, *var2* and *clpr4* seedlings grown on half-strength MS media with 1% sucrose. Bar, 1 mm. Red arrows indicate the first pairs of leaves. (E) Percentages of type a, b and c protoplasts isolated from the first pair of leaves. Data were means ± SD (n = 6). (F) Phenotypes of 20-day-old WT, *var2* and *clpr4* plants grown in soil. Bar, 1 cm. (G) Chloroplast number per guard cell in plants shown in (F). A total of 100 guard cells were analyzed. Statistical analyses were performed with One-Way ANOVA, and significant differences among genotypes were marked with different letters (*P* < 0.05). (H) A general model illustrates the relationship between chloroplast biogenesis, including development and division, and various leaf color phenotypes observed in different mutants. Defects in either chloroplast development, division, or both can lead to distinct leaf phenotypes.

### Interdependence of chloroplast development and division

Chloroplast development and division during leaf development have been suggested to operate independently while mutually compensating to maintain chloroplast biogenesis in response to intracellular and extracellular changes (Pyke, 1999). Similar to the cell cycle, newly divided chloroplasts must grow to a specific size and replicate their DNA before undergoing another round of division (Boffey and Leech, 1982; Ellis and Leech, 1985). Interestingly, impaired chloroplast division appears to have no significant impact on chloroplast development, as chloroplast ultrastructure remains normal in *arc* mutants (Robertson et al., 1995). This interplay between chloroplast development and division is critical for sustaining chloroplast homeostasis during rapid cell proliferation.

The presence of plastid-free cells in *var2* leaves, along with normal cell division, suggests a disruption in the synchronization of chloroplast division with the cell cycle during leaf development. Delayed chloroplast development may hinder chloroplast division by restricting the growth of newly formed chloroplasts. Conversely, defective division exacerbates these developmental delays. This feedback loop likely contributes to the pronounced variegation observed in *var2* leaves.

Based on the available evidence on chloroplast biogenesis, we propose a general model to describe the relationship between chloroplast biogenesis and various leaf phenotypes, including variegation, virescence, and pale green or yellow leaves (Figure 7H). Defects in chloroplast development, as seen in mutants of chlorophyllide *a* oxygenase (*cao*) and divinyl protochlorophyllide 8-vinyl reductase (*pcb2*), result in pale green or yellowish leaves. In contrast, defects in chloroplast division alone typically produce WT-like leaves. However, simultaneous defects in both chloroplast development and division, as in *var2* and *im*, lead to leaf variegation. Accelerated chloroplast division in the context of defective development results in virescent leaves. We previously reported that virescent mutants with reduced plastid gene expression can rescue the *var2* phenotype (Liu et al., 2010; Hu et al., 2015; Ma et al., 2015). Consistent with this, our findings show that chloroplast division is accelerated in virescent mutants such as *clpr4* and *prpl11* (Figures 5C, 7G). This accelerated division may explain the suppression of leaf variegation in virescent mutants. These findings underscore the importance of the interaction between chloroplast development and division in shaping leaf coloration patterns.

In summary, this study advances our understanding of the molecular mechanisms underlying leaf variegation and highlights the coordinated yet independent roles of chloroplast development and division in chloroplast biogenesis. Key conclusions include: (1) Leaf variegation arises from defects in both chloroplast development and division; (2) VAR2 plays a pivotal role in the proplastid-to-chloroplast transition and de-etiolation, likely by promoting chlorophyll biosynthesis; (3) Accelerating chloroplast biogenesis can suppress the variegation phenotype in *var2*. Future research should focus on elucidating the molecular and biochemical mechanisms by which VAR2 regulates chloroplast biogenesis. Understanding these processes will further refine our knowledge of how chloroplast development and division contribute to leaf phenotypic diversity.

## Materials and methods

### Plant materials and growth conditions

*Arabidopsis* seeds used in this study was the Columbia-0 ecotype, and mutants were *var2-1* (Martínez-Zapater, 1993), *pdv1-1* (Miyagishima *et al*., 2006), *pdv2-1* (Miyagishima *et al*., 2006), *arc6-1* (Pyke *et al*., 1994), *parc6-1* (Glynn *et al*., 2009), *prpl11* (Pesaresi *et al*., 2001) and *sot1* (Wu *et al*., 2016). Double mutants, *pdv1 var2*, *pdv2 var2*, *arc6 var2*, *parc6 var2*, *prpl11 var2* and *sot1 var2* were obtained by crossing and identified by sequencing PCR-based fragments. Seeds sterilized with 20% sodium hypochlorite solution were treated at 4 ℃ for 2-3 days, and then sown in half-strength MS plates supplemented with 1% (w/v) sucrose. Seedlings were grown under the long day condition with light intensity of 70-120 µmol m^-2^ s^-1^ at 20 ± 2℃. About 10-day-old seedlings were transplanted into soil for phenotypic observation.

### Plasmid construction and generation of transgenic plants

To generate *PRPL11*-*GFP*, *PDV1*-*FLAG*, and *PDV2*-*FLAG* constructs, *PRPL11*, *PDV1* and *PDV2* CDS sequences were amplified by PCR with the primer pairs listed in Supplemental Table S1, and then cloned into the pENTER vector with an EasyGeno Assembly Cloning Kit (TIANGEN). The pENTER vectors containing target genes were recombined into the binary vector pGWB to get the final vectors. All constructs were confirmed by sequencing. Transgenic plants were screened on half-strength MS plates with antibiotics, and homozygous ones were applied in various assays.

### RNA extraction and quantitative RT-PCR analysis

Total RNA was isolated from the first pair leaves of WT and *var2* plants with an RNA Easy Fast Plant Tissue Kit (TIANGEN), and then reversely transcribed using an PrimeScript™ RT reagent Kit with gDNA Eraser (TaKaRa). The concentrations of cDNA templates were measured by a Nanodrop spectrophotometer before application to qRT-PCR. *ACTIN2* was used as an internal control.

### Green sector size, chlorophyll, and Pchlide measurements

Leaves from 13-d-old and 20-d-old plants were recorded by a stereomicroscope and a scanner. ImageJ software was used to measure the size of green sectors from the first pair of leaves or the whole plant. Percentage of green sector was calculated by the ratio of the green sector area to the whole leaf area.

Chlorophyll was extracted from 0.1 g of rosette leaves with 4 ml of 80% acetone under the dark condition at 4°C on a shaker for 7-8 h until the leaves were completely white, and was measured at OD645 and OD663 using a Nanodrop (Thermo) spectrophotometer. Chlorophyll content was calculated according to the following equations: chlorophyll *a* (μg/ml) = 12.7 x OD663 -2.69 x OD645; chlorophyll *b* (μg/ml) = 22.9 x OD645 - 4.68 x OD663 (Porra *et al*., 1989). Pchlide was extracted from 4-d-old etiolated seedlings with 1.2 ml 80% acetone under the dark condition at 4°C on a shaker overnight, and was measured as previously described (Czarnecki *et al*., 2011).

### Protoplast isolation, confocal microscopy and TEM analysis

Protoplasts were isolated by enzymatic hydrolysis as previously reported (Tan *et al*., 2019), and were observed by a laser confocal fluorescence microscope (Olympus, Tokyo, FV 3000). Plastid numbers were counted using ImageJ software. For FM4-64 dying, 10-day-old seedlings were fully immersed in 10 μM dye solution with a vacuum concentrator for 10 min. The leaf size was recorded by a stereomicroscope, and mesophyll cells were observed by confocal microscope. For propidium iodide (PI) dying, 7-day-old seedlings were stained with the solution (10 μg/ml) for 1.5 min. ImageJ software was used to measure root length and cells. Plastid and chloroplast numbers in guard cells of leaves from 20-day-old *PRPL11-GFP*/WT and *PRPL11-GFP*/*var2* plants grown in soil were recorded by confocal microscope. Data were analyzed using GraphPad Prism 8.0 software. All experiments were repeated at least three times, and similar results were obtained. Statistical analyses were performed with One-Way ANOVA or *t*-test, and significant differences among genotypes were marked with different letters or asterisks (*P* < 0.05).

For TEM observation, SAMs of 3-d-old light-grown seedlings and 4-d-old dark-grown seedlings were fixed and processed as previously described (Zhou *et al*., 2009). Ultrastructure of plastids was observed by a TEM (Tecnai Spirit G2 BioTWIN, FEI, The Kingdom of the Netherlands).

## Funding

This work was supported by the National Natural Science Foundation of China (grant no. 32100191, 32370250); Shanghai Natural Science foundation (21ZR1447100) and Innovation Program of Shanghai Municipal Education Commission (2021-01-07-00-02-E00117).

## Author Contributions

WW, WG and JH designed the study. HZ, WW and WG analyzed correlation between plastid number and cell size. WW, WG, DL, ZZ and MQ made transgenic plants and mutants. WW, WG, ZY and JH conducted confocal microscopy and TEM analyses. WW, WG and JH wrote the manuscript. All authors read and approved the manuscript.

## Declaration of interests

The authors declare no competing interests.

## Supplementary information

**Table S1. Primers used in the experiments.**

**Figure S1. Overexpression of RPL11-GFP has no obvious effect on leaf variegation of *var2*.**

**Figure S2. Quantitative PCR analysis of expression levels of chloroplast division-related genes.**

**Figure S3. Identification of double mutants, *var2* with *arc6*, *parc6*, *pdv1* or *pdv2* via semi-quantitative RT-PCR.**

**Figure S4. Semi-quantitative RT-PCR analysis of transgenic plants overexpressing *PDV1* and *PDV2* in *var2*.**

**Figure S5. Early chloroplast biogenesis is critical for suppression of leaf variegation.**

**Figure S6. *COP1* mutations can rescue variegated leaf phenotype of *var2* under various light/dark cycles.**

## Notes

### Competing Interest Statement

The authors have declared no competing interest.

## References

Aluru MR, Yu F, Fu A, Rodermel S. 2006. Arabidopsis variegation mutants: new insights into chloroplast biogenesis. J Exp Bot 57, 1871–1881.

Asano T, Yoshioka Y, Kurei S, Sakamoto W, Machida Y. 2004. A mutation of the CRUMPLED LEAF gene that encodes a protein localized in the outer envelope membrane of plastids affects the pattern of cell division, cell differentiation, and plastid division in Arabidopsis. Plant J 38, 448–459.

Birky CW, Jr. 1983. The partitioning of cytoplasmic organelles at cell division. Int Rev Cytol Suppl 15, 49–89.

Boffey SA, Leech RM. 1982. Chloroplast DNA levels and the control of chloroplast division in light-grown wheat leaves. Plant Physiol 69, 1387–1391.

Charuvi D, Kiss V, Nevo R, Shimoni E, Adam Z, Reich Z. 2012. Gain and loss of photosynthetic membranes during plastid differentiation in the shoot apex of Arabidopsis. Plant Cell 24, 1143–1157.

Chen L, Sun B, Gao W, Zhang QY, Yuan H, Zhang M. 2018. MCD1 Associates with FtsZ Filaments via the Membrane-Tethering Protein ARC6 to Guide Chloroplast Division. Plant Cell 30, 1807–1823.

Chen Y, Asano T, Fujiwara MT, Yoshida S, Machida Y, Yoshioka Y. 2009. Plant cells without detectable plastids are generated in the crumpled leaf mutant of Arabidopsis thaliana. Plant Cell Physiol 50, 956–969.

Czarnecki O, Peter E, Grimm B. 2011. Methods for analysis of photosynthetic pigments and steady-state levels of intermediates of tetrapyrrole biosynthesis. Methods Mol Biol 775, 357–385.

Dogra V, Duan J, Lee KP, Kim C. 2019. Impaired PSII proteostasis triggers a UPR-like response in the var2 mutant of Arabidopsis. J Exp Bot 70, 3075–3088.

Ellis JR, Leech RM. 1985. Cell size and chloroplast size in relation to chloroplast replication in light-grown wheat leaves. Planta 165, 120–125.

Fang J, Li B, Chen LJ, Dogra V, Luo S, Wu W, Wang P, Hwang I, Li HM, Kim C. 2022. TIC236 gain-of-function mutations unveil the link between plastid division and plastid protein import. Proc Natl Acad Sci U S A 119, e2123353119.

Gao H, Kadirjan-Kalbach D, Froehlich JE, Osteryoung KW. 2003. ARC5, a cytosolic dynamin-like protein from plants, is part of the chloroplast division machinery. Proc Natl Acad Sci U S A 100, 4328–4333.

Glynn JM, Froehlich JE, Osteryoung KW. 2008. Arabidopsis ARC6 coordinates the division machineries of the inner and outer chloroplast membranes through interaction with PDV2 in the intermembrane space. Plant Cell 20, 2460–2470.

Glynn JM, Yang Y, Vitha S, Schmitz AJ, Hemmes M, Miyagishima SY, Osteryoung KW. 2009. PARC6, a novel chloroplast division factor, influences FtsZ assembly and is required for recruitment of PDV1 during chloroplast division in Arabidopsis. Plant J 59, 700–711.

Han X, Huang X, Deng XW. 2020. The Photomorphogenic Central Repressor COP1: Conservation and Functional Diversification during Evolution. Plant Commun 1, 100044.

Haswell ES, Meyerowitz EM. 2006. MscS-like proteins control plastid size and shape in Arabidopsis thaliana. Curr Biol 16, 1–11.

He H, Xie W, Liang Z, Wu H, Bai M. 2021. The expansion of mesophyll cells is coordinated with the division of chloroplasts in diploid and tetraploid Arabidopsis thaliana. Planta 253, 64.

Hu F, Zhu Y, Wu W, Xie Y, Huang J. 2015. Leaf Variegation of Thylakoid Formation1 Is Suppressed by Mutations of Specific σ-Factors in Arabidopsis. Plant Physiol 168, 1066–1075.

Hudik E, Yoshioka Y, Domenichini S, Bourge M, Soubigout-Taconnat L, Mazubert C, Yi D, Bujaldon S, Hayashi H, De Veylder L, Bergounioux C, Benhamed M, Raynaud C. 2014. Chloroplast dysfunction causes multiple defects in cell cycle progression in the Arabidopsis crumpled leaf mutant. Plant Physiol 166, 152–167.

Izumi M, Ishida H, Nakamura S, Hidema J. 2017. Entire Photodamaged Chloroplasts Are Transported to the Central Vacuole by Autophagy. Plant Cell 29, 377–394.

Jarvis P, López-Juez E. 2013. Biogenesis and homeostasis of chloroplasts and other plastids. Nat Rev Mol Cell Biol 14, 787–802.

Kato Y, Hyodo K, Sakamoto W. 2018. The Photosystem II Repair Cycle Requires FtsH Turnover through the EngA GTPase. Plant Physiol 178, 596–611.

Kato Y, Kuroda H, Ozawa SI, Saito K, Dogra V, Scholz M, Zhang G, de Vitry C, Ishikita H, Kim C, Hippler M, Takahashi Y, Sakamoto W. 2023. Characterization of tryptophan oxidation affecting D1 degradation by FtsH in the photosystem II quality control of chloroplasts. Elife 12.

Kato Y, Miura E, Ido K, Ifuku K, Sakamoto W. 2009. The variegated mutants lacking chloroplastic FtsHs are defective in D1 degradation and accumulate reactive oxygen species. Plant Physiol 151, 1790–1801.

Kato Y, Miura E, Matsushima R, Sakamoto W. 2007. White leaf sectors in yellow variegated2 are formed by viable cells with undifferentiated plastids. Plant Physiol 144, 952–960.

Liu X, Yu F, Rodermel S. 2010. Arabidopsis chloroplast FtsH, var2 and suppressors of var2 leaf variegation: a review. J Integr Plant Biol 52, 750–761.

Ma Z, Wu W, Huang W, Huang J. 2015. Down-regulation of specific plastid ribosomal proteins suppresses thf1 leaf variegation, implying a role of THF1 in plastid gene expression. Photosynth Res 126, 301–310.

Martínez-Zapater JM. 1993. Genetic Analysis of Variegated Mutants in Arabidopsis. Journal of Heredity 84, 138–140.

Miura E, Kato Y, Matsushima R, Albrecht V, Laalami S, Sakamoto W. 2007. The balance between protein synthesis and degradation in chloroplasts determines leaf variegation in Arabidopsis yellow variegated mutants. Plant Cell 19, 1313–1328.

Miyagishima SY. 2011. Mechanism of plastid division: from a bacterium to an organelle. Plant Physiol 155, 1533–1544.

Miyagishima SY, Froehlich JE, Osteryoung KW. 2006. PDV1 and PDV2 mediate recruitment of the dynamin-related protein ARC5 to the plastid division site. Plant Cell 18, 2517–2530.

Miyagishima SY, Kabeya Y. 2010. Chloroplast division: squeezing the photosynthetic captive. Curr Opin Microbiol 13, 738–746.

Niwa Y, Kato T, Tabata S, Seki M, Kobayashi M, Shinozaki K, Moriyasu Y. 2004. Disposal of chloroplasts with abnormal function into the vacuole in Arabidopsis thaliana cotyledon cells. Protoplasma 223, 229–232.

Osteryoung KW, Pyke KA. 2014. Division and dynamic morphology of plastids. Annu Rev Plant Biol 65, 443–472.

Pedroza-Garcia JA, Domenichini S, Bergounioux C, Benhamed M, Raynaud C. 2016. Chloroplasts around the plant cell cycle. Curr Opin Plant Biol 34, 107–113.

Pesaresi P, Varotto C, Meurer J, Jahns P, Salamini F, Leister D. 2001. Knock-out of the plastid ribosomal protein L11 in Arabidopsis: effects on mRNA translation and photosynthesis. Plant J 27, 179–189.

Porra RJ, Thompson WA, Kriedemann PE. 1989. Determination of accurate extinction coefficients and simultaneous equations for assaying chlorophylls a and b extracted with four different solvents: verification of the concentration of chlorophyll standards by atomic absorption spectroscopy. Biochimica et Biophysica Acta (BBA) - Bioenergetics 975, 384–394.

Pyke K. 1997. The genetic control of plastid division in higher plants. Am J Bot 84, 1017.

Pyke KA. 1999. Plastid division and development. Plant Cell 11, 549–556.

Pyke KA, Rutherford SM, Robertson EJ, Leech RM. 1994. arc6, A Fertile Arabidopsis Mutant with Only Two Mesophyll Cell Chloroplasts. Plant Physiol 106, 1169–1177.

Robertson EJ, Pyke KA, Leech RM. 1995. arc6, an extreme chloroplast division mutant of Arabidopsis also alters proplastid proliferation and morphology in shoot and root apices. J Cell Sci 108 ( Pt 9), 2937–2944.

Rosso D, Bode R, Li W, Krol M, Saccon D, Wang S, Schillaci LA, Rodermel SR, Maxwell DP, Hüner NP. 2009. Photosynthetic redox imbalance governs leaf sectoring in the Arabidopsis thaliana variegation mutants immutans, spotty, var1, and var2. Plant Cell 21, 3473–3492.

Sakamoto W. 2003. Leaf-variegated mutations and their responsible genes in Arabidopsis thaliana. Genes Genet Syst 78, 1–9.

Sakamoto W. 2006. Protein degradation machineries in plastids. Annu Rev Plant Biol 57, 599–621.

Sakamoto W, Miyagishima SY, Jarvis P. 2008. Chloroplast biogenesis: control of plastid development, protein import, division and inheritance. Arabidopsis Book 6, e0110.

Sakamoto W, Uno Y, Zhang Q, Miura E, Kato Y, Sodmergen. 2009. Arrested differentiation of proplastids into chloroplasts in variegated leaves characterized by plastid ultrastructure and nucleoid morphology. Plant Cell Physiol 50, 2069–2083.

Schwab R, Ossowski S, Riester M, Warthmann N, Weigel D. 2006. Highly specific gene silencing by artificial microRNAs in Arabidopsis. Plant Cell 18, 1121–1133.

Sheahan MB, Rose RJ, McCurdy DW. 2004. Organelle inheritance in plant cell division: the actin cytoskeleton is required for unbiased inheritance of chloroplasts, mitochondria and endoplasmic reticulum in dividing protoplasts. Plant J 37, 379–390.

Sumiya N, Fujiwara T, Era A, Miyagishima SY. 2016. Chloroplast division checkpoint in eukaryotic algae. Proc Natl Acad Sci U S A 113, E7629–e7638.

Sun B, Zhang QY, Yuan H, Gao W, Han B, Zhang M. 2020. PDV1 and PDV2 Differentially Affect Remodeling and Assembly of the Chloroplast DRP5B Ring. Plant Physiol 182, 1966–1978.

Tan H, Man C, Xie Y, Yan J, Chu J, Huang J. (2019). A crucial role of GA-regulated flavonol biosynthesis in root growth of Arabidopsis. Molecular plant 12, 521–537.

Tsukaya H. 2002. Leaf development. Arabidopsis Book 1, e0072.

Tsukaya H. 2013. Leaf development. Arabidopsis Book 11, e0163.

Wang W, Li J, Sun Q, Yu X, Zhang W, Jia N, An C, Li Y, Dong Y, Han F, Chang N, Liu X, Zhu Z, Yu Y, Fan S, Yang M, Luo SZ, Gao H, Feng Y. 2017. Structural insights into the coordination of plastid division by the ARC6-PDV2 complex. Nat Plants 3, 17011.

Wilson ME, Jensen GS, Haswell ES. 2011. Two mechanosensitive channel homologs influence division ring placement in Arabidopsis chloroplasts. Plant Cell 23, 2939–2949.

Wu W, Liu S, Ruwe H, Zhang D, Melonek J, Zhu Y, Hu X, Gusewski S, Yin P, Small ID, Howell KA, Huang J. 2016. SOT1, a pentatricopeptide repeat protein with a small MutS-related domain, is required for correct processing of plastid 23S-4.5S rRNA precursors in Arabidopsis thaliana. Plant J 85, 607–621.

Wu W, Zhu Y, Ma Z, Sun Y, Quan Q, Li P, Hu P, Shi T, Lo C, Chu IK, Huang J. 2013. Proteomic evidence for genetic epistasis: ClpR4 mutations switch leaf variegation to virescence in Arabidopsis. Plant J 76, 943–956.

Yadav D, Zemach H, Belausov E, Charuvi D. 2019. Initial proplastid-to-chloroplast differentiation in the developing vegetative shoot apical meristem of Arabidopsis. Biochem Biophys Res Commun 519, 391–395.

Yu F, Fu A, Aluru M, Park S, Xu Y, Liu H, Liu X, Foudree A, Nambogga M, Rodermel S. 2007. Variegation mutants and mechanisms of chloroplast biogenesis. Plant Cell Environ 30, 350–365.

Yu F, Park S, Rodermel SR. 2004. The Arabidopsis FtsH metalloprotease gene family: interchangeability of subunits in chloroplast oligomeric complexes. Plant J 37, 864–876.

Zhang M, Chen C, Froehlich JE, TerBush AD, Osteryoung KW. 2016. Roles of Arabidopsis PARC6 in Coordination of the Chloroplast Division Complex and Negative Regulation of FtsZ Assembly. Plant Physiol 170, 250–262.

Zhou W, Cheng Y, Yap A, Chateigner-Boutin AL, Delannoy E, Hammani K, Small I, Huang J. 2009. The Arabidopsis gene YS1 encoding a DYW protein is required for editing of rpoB transcripts and the rapid development of chloroplasts during early growth. Plant J 58, 82–96.

