## Supplemental Data for "An unrecognized and crucial role of chloroplast division in leaf variegation in *Arabidopsis thaliana*"

**Table S1 Primers used in the experiments.**

| Names | Sequences(5'→3') |
|---|---|
| *PDV1*-RT-F | GCGATGAGATTCACTTGATTTC |
| *PDV1*-RT-R | CGTATGTTTTCCTTTGTTGGC |
| *PDV2*-RT-F | TTTGCGTATTCGTGATGCTCTTG |
| *PDV2*-RT-R | CCATTTGACTCTGTTGTAGCGAA |
| *ARC6*-RT-F | CTGAGATAGTTCTTCGGGTTGGT |
| *ARC6*-RT-R | AATCTCCATAGCCATTACCTTAGC |
| *PARC6*-RT-F | CGAGAAGGACGGAGGAGGTTGAA |
| *PARC6*-RT-R | CTCAGCAAGTGCCATAGATAAGA |
| *ACTIN2*-RT-F | TCTTCTTCCGCTCTTTCTTTCC |
| *ACTIN2*-RT-R | TCTTACAATTTCCCGCTCTGC |
| *ARC6*-q-F | ATGGCGCTTGCGTTTCTCGATG |
| *ARC6*-q-R | AAGGCTACTTGCTCCTTCCTCCTG |
| *PARC6*-q-F | TGATGAATCCATGCTTGTCCAGTG |
| *PARC6*-q-R | CAGGATTTCGCCTCTGCTGTTTG |
| *PDV1*-q-F | GTTTTCATCAAAGGATTCGCCGTC |
| *PDV1*-q-R | CTGTTCGCTGCTGTGTATGTATAG |
| *PDV2*-q-F | AACGGCTTTTGCGTATTCGTGATG |
| *PDV2*-q-R | CTTCTCAAGCAACATTTTCCTACT |
| *ACTIN2*-q-F | CACTTGCACCAAGCAGCATGAAGA |
| *ACTIN2*-q-R | AATGGAACCACCGATCCAGACACT |

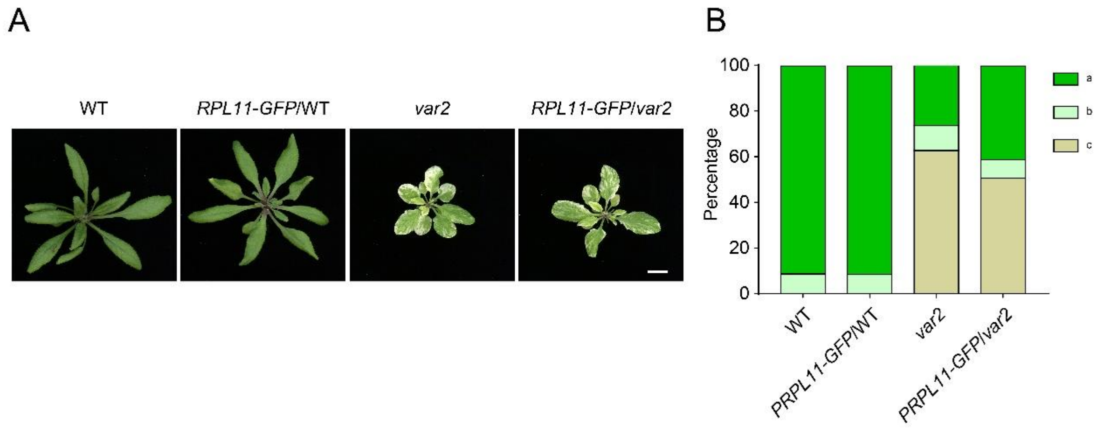


**Figure S1. Overexpression of *RPL11-GFP* has no obvious effect on leaf variegation of *var2*.**

(A) Phenotypes of 20-day-old WT, *RPL11-GFP*/WT, *var2*, and *RPL11-GFP*/*var2* plants grown in soil in the 16 hr light/8 hr dark cycle. Bar, 1 cm. (B) Percentage of type a, b and c cells isolated from plants shown in (A). More than 300 protoplasts were analyzed.

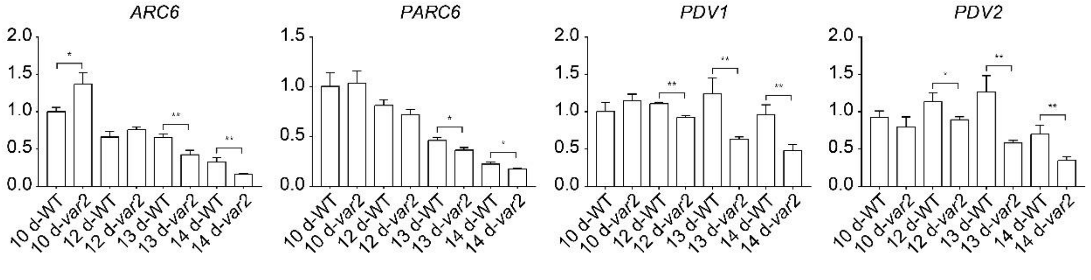


**Figure S2. Quantitative PCR analysis of expression levels of chloroplast division-related genes.**

Total RNA was extracted from the first pair of leaves from 10, 12, 13, and 14-day-old WT and *var2* seedlings grown on half-strength MS media with 1% sucrose. The expression levels of *ARC6*, *PARC6*, *PDV1*, and *PDV2* were analyzed. Data were means ± SD (n = 3). Significant difference between WT and *var2* was labeled with one star ($P < 0.05$) and two stars ($P < 0.01$) (student *t*-test). *ACTIN2* was used as an internal control.

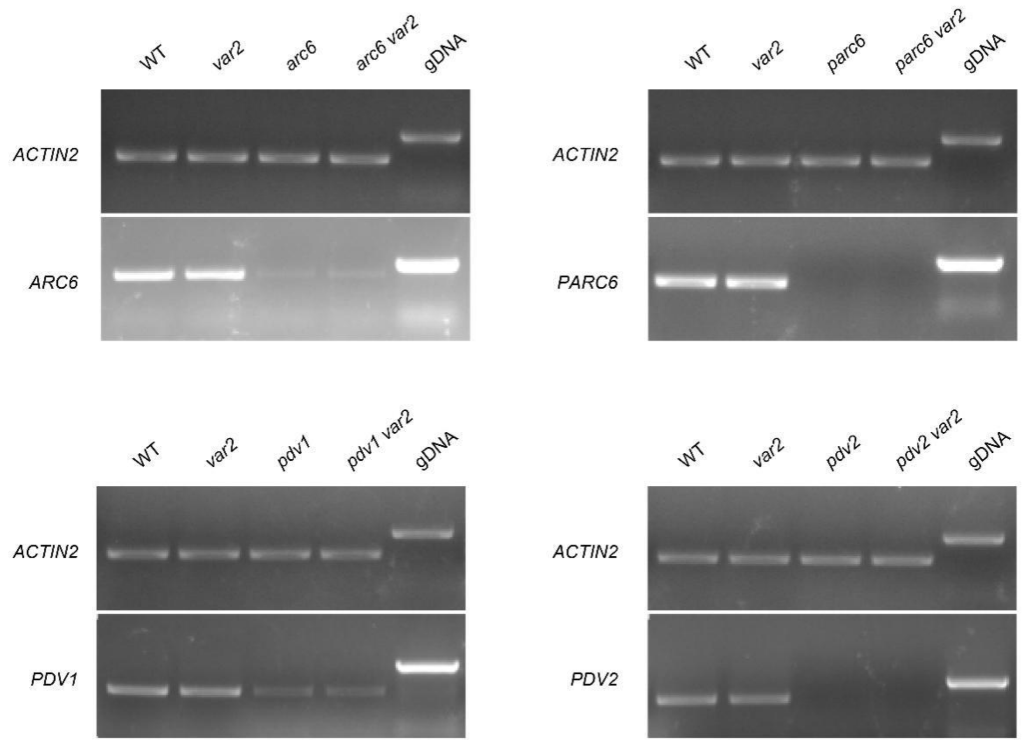


**Figure S3. RT-PCR analysis of double mutants, *var2 arc6*, *var2 parc6*, *var2 pdv1* and *var2 pdv2*.**

Total RNA was isolated from the leaves of 10-day-old seedlings grown on half-strength MS media in the 16 hr light/8 hr dark cycle. *ACTIN2* was used as a control. It should be pointed out that mRNA detected in genetic backgrounds containing *arc6* and *pdv1* are knockout mutants since *arc6* and *pdv1* are a single-base mutation, leading to premature termination in protein translation.

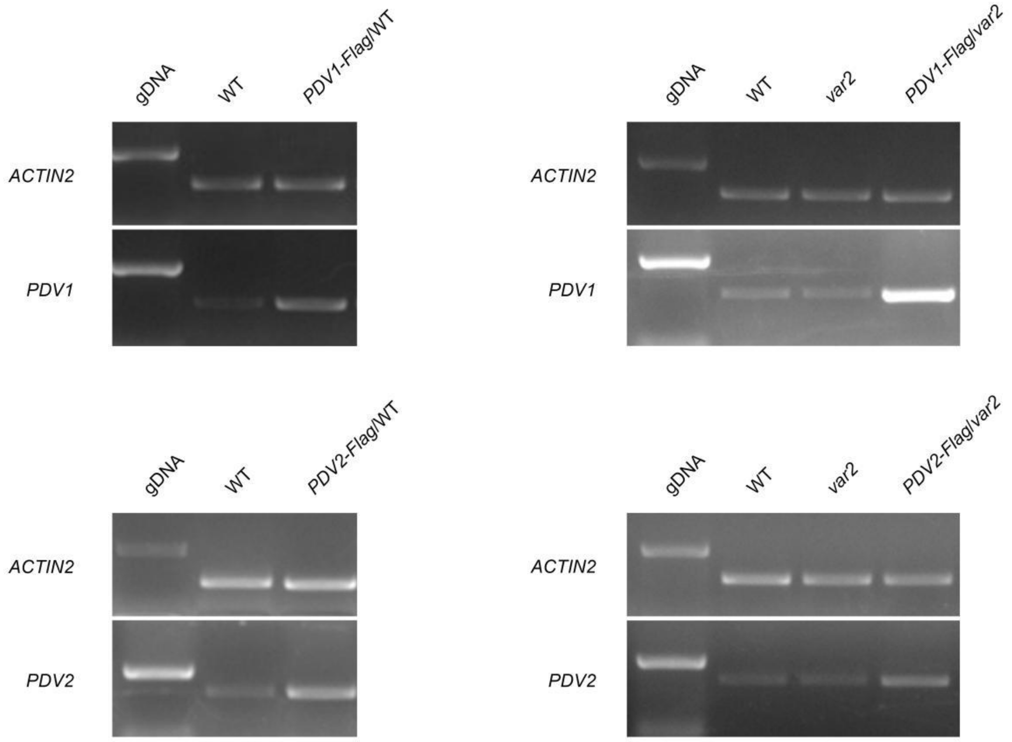


**Figure S4. RT-PCR analysis of transgenic plants overexpressing *PDV1* and *PDV2* in *var2*.**

Total RNA was isolated from leaves of 20-day-old WT, *var2*, *PDV1-Flag*/WT, *PDV2-Flag*/WT, *PDV1-Flag*/*var2*, and *PDV1-Flag*/*var2* plants grown in soil in the 16 hr light/8 hr dark cycle. *ACTIN2* was used as a loading control.

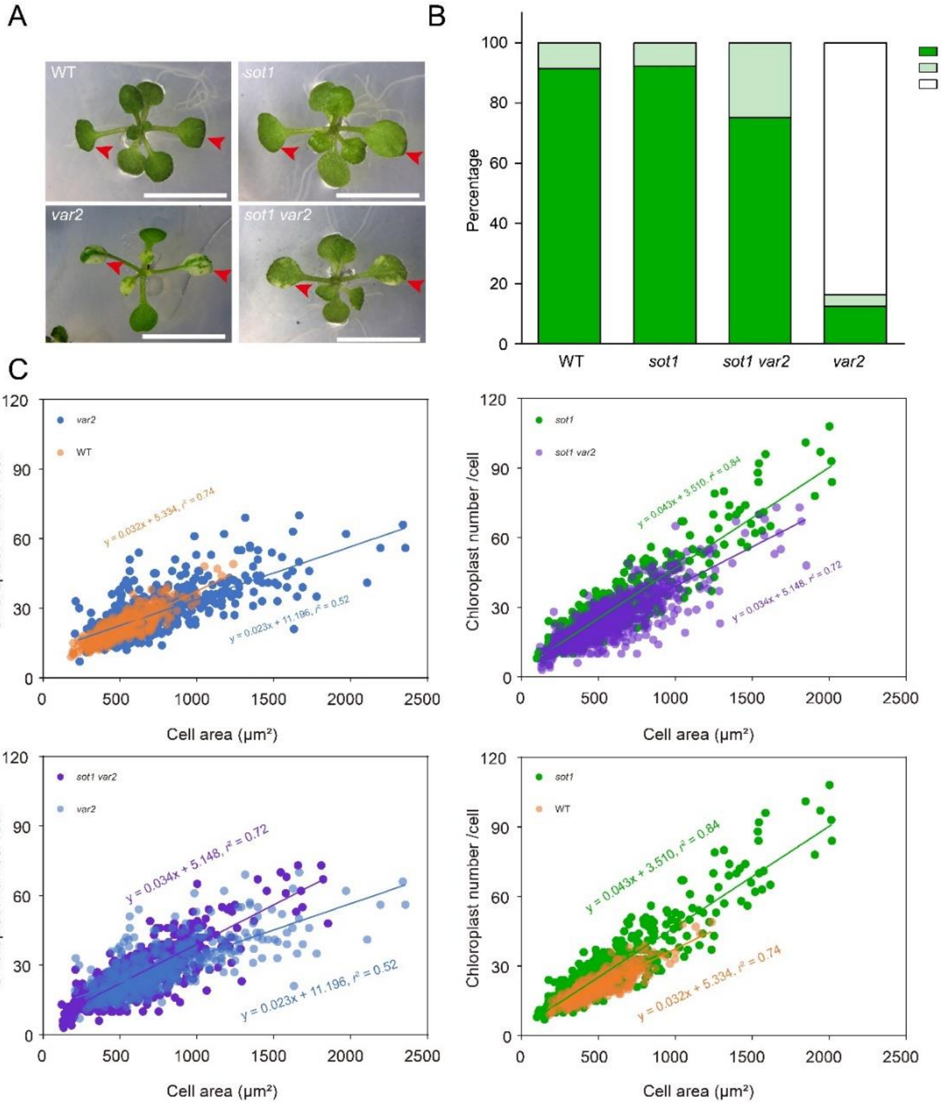


**Figure S5. Early chloroplast biogenesis is critical for suppression of leaf variegation.**

(A) Phenotypes of 13-day-old WT, *var2*, *sot1* and *sot1 var2* seedlings cultured on half-strength MS media. Bars, 0.5 cm. Red arrows indicate the first pair of true leaves. (B) Percentage of type a, b, or c cells. More than 400 protoplasts isolated from the first pair of leaves of the plants shown in A were analyzed. (C) Correlation analysis between chloroplast number and cell area in type a cells. More than 500 cells were analyzed for each sample. The $r^2$ values of the best-fit lines are 0.74 (WT), 0.52 (*var2*), 0.84 (*sot1*), and 0.72 (*sot1 var2*). The $k$ values are 0.032 (WT), 0.023 (*var2*), 0.043 (*sot1*), and 0.034 (*sot1 var2*).

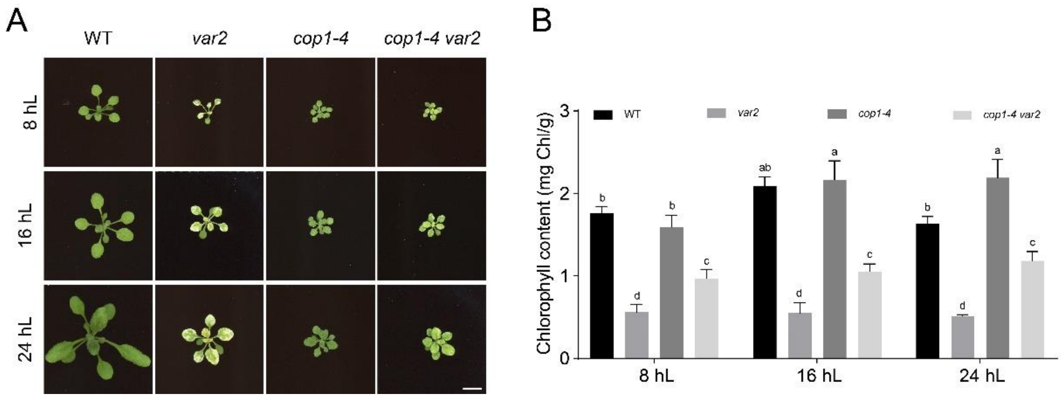


**Figure S6. *COP1* mutations rescue variegated leaf phenotype of *var2* under various light/dark cycles.**

(A) Phenotypes of 10-day-old WT, *var2*, *cop1-4*, and *cop1-4 var2* plants grown in soil under different light cycle conditions. Bar, 1 cm. (B) Chlorophyll contents of plants shown in (A). The data were shown means ± SD (n = 3). Significant differences among different genotypes and different conditions (One-Way ANOVA, $P < 0.05$) were marked with different letters.
